# Postinspiratory complex acts as a gating mechanism regulating swallow-breathing coordination and other laryngeal behaviors

**DOI:** 10.1101/2023.01.18.524513

**Authors:** Alyssa Huff, Marlusa Karlen-Amarante, Luiz Marcelo Oliveira, Jan Marino Ramirez

## Abstract

Breathing needs to be tightly coordinated with upper airway behaviors, such as swallowing. Discoordination leads to aspiration pneumonia, the leading cause of death in neurodegenerative diseases. Here we study the role of the postinspiratory complex, (PiCo) in coordinating breathing and swallowing. Using optogenetic approaches in freely breathing-anesthetized ChATcre, Vglut2cre and co-transmission of ChATcre/Vglut2FlpO mice reveals this small brainstem microcircuit acts as a central gating mechanism for airway protective behaviors. Activation of PiCo during inspiration or the beginning of postinspiration triggers swallow behavior, while there is a higher probability for stimulating laryngeal activation when activated further into expiration, suggesting PiCo’s role in swallow-breathing coordination. PiCo triggers consistent swallow behavior and preserves physiologic swallow motor sequence, while stimulates laryngeal activation variable to stimulation duration. Sufficient bilateral PiCo activation is necessary for gating function since activation of only a few PiCo neurons or unilateral activation leads to blurred behavioral response. Viral tracing experiments reveal projections from the caudal nucleus of the solitary tract (cNTS), the presumed swallow pattern generator (SPG), to PiCo and vice versa. However, PiCo does not directly connect to laryngeal muscles. Investigating PiCo’s role in swallow and laryngeal coordination will aid in understanding discoordination in breathing and neurological diseases.

## Introduction

The discovery of the preBötzinger complex (preBötC) in the brainstem 30 years ago triggered a wave of mechanistic studies aimed at understanding the neuronal determinants that drive inhalation. By contrast, the mechanistic understanding of exhalation lacks far behind. Indeed, often called “passive” expiration this term suggests that expiration is primarily driven by mechanical recoil forces of the lung and may occur without neuronal control. Far from the truth, expiration is complex involving the neuronal control of multiple muscles, and the exquisite coordinated valving of laryngeal and pharyngeal control with other behaviors such as vocalization, coughing or swallowing.

The complexity of exhalation is partly reflected in the fact that it can be subdivided into different phases – exhalation begins with postinspiration or E1 phase, followed by late expiration or the E2 phase (D. W. Richter & Smith, 2014). But, there is also “active expiration” which is associated with the conditional activation of intercostal and abdominal muscles that are recruited during high metabolic demand to actively exhale (Abdala et al., 2009; Flor, Barnett, Karlen-Amarante, Molkov, & Zoccal, 2020; Molkov et al., 2011). Over the years and throughout the field of breathing expiration has been used in multiple fashions. Postinspiration was originally defined in 1937 by Gesell and White (Gesell & White, 1938) calling it an “after discharge” of the diaphragm, similar to “yield” defined by Huff et al (Huff A, 2020a). Believed to not be remnant activity of the preceding inspiratory discharge but a separate activity essential for rhythmogenesis (D. Richter, 1982). In 1973 Gautier, Remmers and Bartlett documented the functional importance of postinspiration as an expiratory braking mechanism, suggesting this action would be more suitable for laryngeal control due to its fine motor control (Bartlett Jr, Remmers, & Gautier, 1973; Gautier, Remmers, & Bartlett Jr, 1973), similar to Dutschmann et al (Dutschmann & Dick, 2012) defined as early expiration or E1. While, Bautista introduces postinspiration as interchangeable with early expiration (Bautista & Dutschmann, 2014).

In 2016, Anderson et al. described PiCo as a heterogenous population of interneurons, located within the intermediate reticular nucleus (IRt), that uniquely co-expresses both glutamate and acetylcholine and was found to be both sufficient and necessary for generating this postinspiratory phase of breathing (Anderson et al., 2016). It was hypothesized this complex may be involved in various postinspiratory behaviors such as swallowing and vocalization (Anderson et al., 2016). Since, work in the rodent has shown this region serves as a premotor relay within the IRt that integrates postinspiratory motor outputs and other non-respiratory central pattern generators (CPGs) such as swallowing, crying, lapping, whisking (Ain Summan Toor et al., 2019; Dempsey et al., 2021; Moore et al., 2013; Moore, Kleinfeld, & Wang, 2014; Pitts, Huff, Reed, Iceman, & Mellen, 2021), suggesting that PiCo may serve as a hub, acting as a gate, for various laryngeal postinspiratory behaviors.

In locomotion, sensory information about limb position gates a motor response elicited by sensory stimulus resulting in different motor output depending on the phase of locomotion the sensory stimuli appears (Grillner, 2006). For example, identical stimulus applied to the paw will activate flexors during the swing phase of movement, but extensors during the stance phase, mediated at an interneuronal, premotoneuron level (Andersson, Forssberg, Grillner, & Lindquist, 1978; Forssberg, Grillner, & Rossignol, 1975; Forssberg, Grillner, & Rossignol, 1977; Grillner, 2006).

In this study we aimed to further explore the role of PiCo in the coordination of breathing, swallowing and laryngeal activation, a behavior important for respiration (Bartlett, 1986; Bartlett Jr, 1989; Dutschmann & Dick, 2012) and during swallow provides airway protection. Using optogenetic techniques, phase specific activation of *ChAT, Vglut2*, and cotransmission of *ChAT/Vglut2* neurons at the level of PiCo in a spontaneously breathing anesthetized *in vivo* preparation, resulted in two airway protective behaviors: swallow and laryngeal activation. We hypothesize that PiCo acts as a gating mechanism for airway protective behaviors.

## Results

### Optogenetic stimulation of PiCo neurons gates swallow and laryngeal activation in a phase specific manner

#### Optogenetic stimulation of ChAT neurons at PiCo

Activation of ChAT neurons in PiCo leads to laryngeal muscle activation or a swallow dependent on the timing within the respiratory cycle. A two-way ANOVA revealed swallow is triggered with a significantly higher probability when ChAT neurons are activated within the first 10% (*p*= 0.02) of the respiratory cycle. However, there is a significantly higher probability laryngeal muscle activation will occur when ChAT neurons are activated within 70% (*p*= 0.04) to 90% (*p*= 0.005) of the respiratory cycle (Fig. 1A).

**Figure 1.**
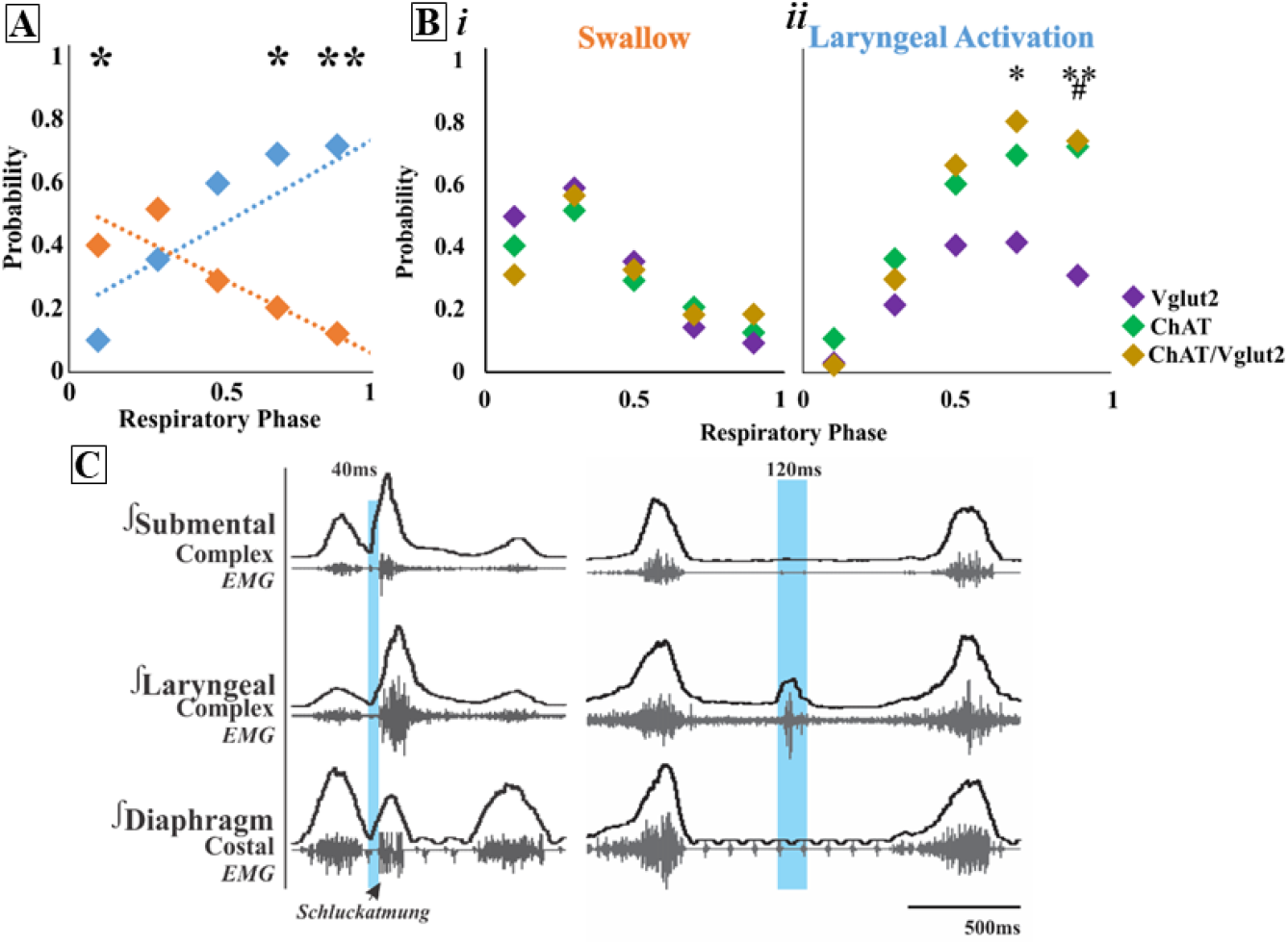
Optogenetic stimulation of PiCo neurons gates swallow and laryngeal activation in a phase specific manner. A) Scatter plot of probability of triggering a swallow (orange) or laryngeal activation (blue) across the respiratory phase (0 start of inspiration, 1 start of next inspiration) in ChAT mice. * Indicates significant difference between probability of evoking a swallow or laryngeal activation within the first 10% (*p* = 0.02), 70% (*p*= 0.04), and **90% (*p*=0.005) of the respiratory cycle. Bi) Scatter plot of the probability of triggering a swallow shows no difference between Vglut2 (purple), ChAT Ai32 (green) and ChAT/Vglut2 (gold) mice. Bii) There is no change in probability of stimulating laryngeal activation between ChAT and ChAT/Vglut2 mice, however there is a significant difference between Vglut2 and *ChAT/Vglut2 mice at 70% (*p*= 0.04) and 90% (*p*= 0.02) of the respiratory cycle and Vglut2 and #ChAT mice at 90%(*p*= 0.008) of the respiratory cycle. C) Representative traces of PiCo triggered swallow on the left showing swallow motor sequence of submental and laryngeal activation, plus swallow related diaphragm activation known as *Schluckatmung*. Characterization of laryngeal activation on the right showing only the laryngeal complex is activated in response to the laser in blue.

#### Optogenetic stimulation of Vglut2 neurons at PiCo

There is also a significantly higher probability a swallow will be triggered when Vglut2 neurons are stimulated in PiCo within the first 10% (*p*<0.0001) to 30% (*p*= 0.002) of the respiratory cycle (Fig. 1Bi), while laryngeal activation will occur with a significantly higher probability when Vglut2 neurons are activated within 70% (*p*= 0.04) of the respiratory cycle (Fig. 1Bii).

#### Optogenetic stimulation of ChAT/Vglut2 neurons at PiCo

To specifically stimulate PiCo neurons, we used double conditioned mice expressing cre in ChAT cells and FlpO in Vglut2 cells. We then injected the pAAV-hSyn Con/Fon hChR2(H134R)-EYFP vector into PiCo, resulting in expression of channelrhodopsin in neurons that only co-express ChAT and Vglut2, herein will be referred to as ChAT/Vglut2. There is a significantly higher probability laryngeal activation will be stimulated when PiCo neurons are activated within 70% (*p*= 0.04) of the respiratory cycle.

When comparing the probability of triggering a swallow between ChAT, Vglut2, and ChAT/Vglut2 mice there is no significant difference across the respiratory cycle (Fig. 1Bi). However, the probability of triggering laryngeal activation in Vglut2 mice compared to ChAT/Vglut2 mice is significantly lower at 70% (*p*= 0.04) and 90% (*p*= 0.02) of the respiratory cycle (Fig. 1Bii). Also, the probability of triggering laryngeal activation in Vglut2 mice compared to ChAT mice is significantly lower at 90% (*p*= 0.008) of the respiratory cycle (Fig. 1Bii).

When evaluating the phase shift plots, we divided PiCo stimulated responses into swallow and non-swallow (Fig. 2C). Non-swallow included PiCo activation that resulted in both laryngeal activation and no response. We found that in all genetic mouselines studied, laser pulse duration did not affect respiratory rhythm reset in either swallow or non-swallow responses, allowing us to group all laser pulse durations as one. R values were calculated for each genetic mouse type and response. In figure 2A, swallow stimulated by ChAT r = 0.77 (*p*< 0.0001), Vglut2 r = 0.33 (*p*< 0.0001), and ChAT/Vglut2 r = 0.75 (*p*< 0.0001). Non-swallows stimulated by ChAT r = 0.45 (*p*< 0.0001), Vglut2 r = 0.29 (*p*< 0.0001), and ChAT/Vglut2 r = 0.18 (*p*= 0.0001). In figure 2B the line of best fit was calculated for both responses and genetic type. Swallows stimulated by ChAT R^2^ = 0.59, Vglut2 R^2^= 0.11, and ChAT/Vglut2 R^2^ = 0.57. Non-swallows stimulated by ChAT R^2^ = 0.20, Vglut2 R^2^ = 0.09, and ChAT/Vglut2 R^2^ = 0.03.

**Figure 2.**
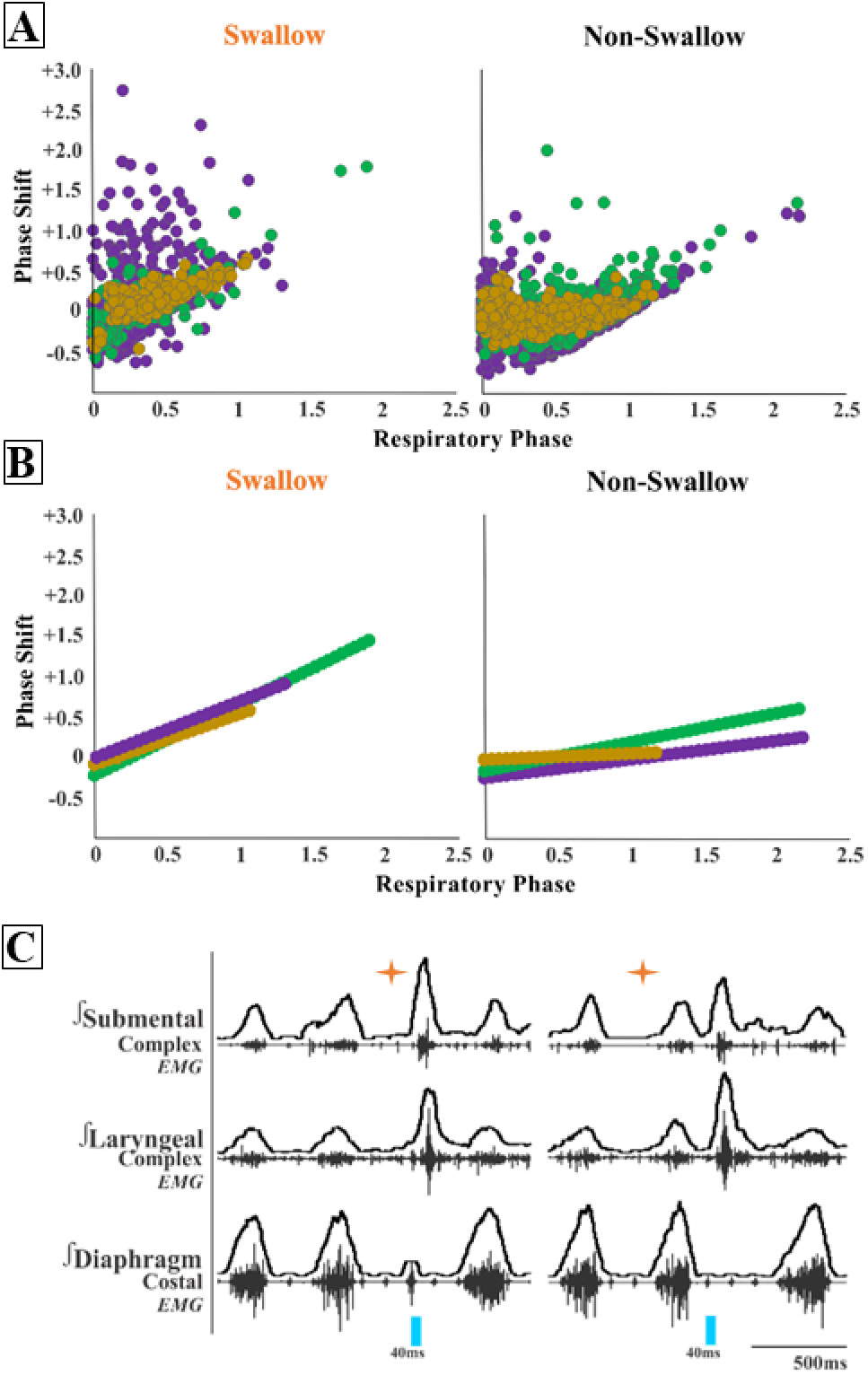
PiCo triggered swallows resets the respiratory rhythm, while non swallows have minimal affect. Respiratory phase shifts plots were divided into two groups: swallow, PiCo laser activation that triggered a swallow, or non-swallow, PiCo activation that resulted in laryngeal activation or no response. A) Individual laser response in ChAT/Vglut2 (gold), ChAT (green), and Vglut2 (purple) and B) line of best fit from the above graphs. C) Representative traces of two examples of swallow (orange star) response on respiratory cycle. On the left, PiCo triggered swallow inhibits inspiration resulting in an earlier onset of the next inspiratory breath, and on the right a delay in the next inspiration.

### PiCo stimulation triggers swallow behavior while activates laryngeal activity

We found that regardless of laser pulse duration, ranging from 40ms to 200ms, swallows were triggered in an all-or-none manner and had an average duration of 116 ± 20ms in the ChAT/Vglut2 mice (Fig. 3A). Laser pulse duration has no effect on PiCo triggered swallow response. By contrast, PiCo, activated laryngeal activity in a gradual manner. As laser pulse duration increased, laryngeal duration increased in an on-off fashion where responses to 40ms pulses were significantly shorter than laryngeal activity stimulated with 200ms pulses (*p*<0.0001) (Fig. 3A). This was also true in both the ChAT (*p*<0.02) and Vglut2 (*p*<0.004) mice (Table S1).

**Figure 3.**
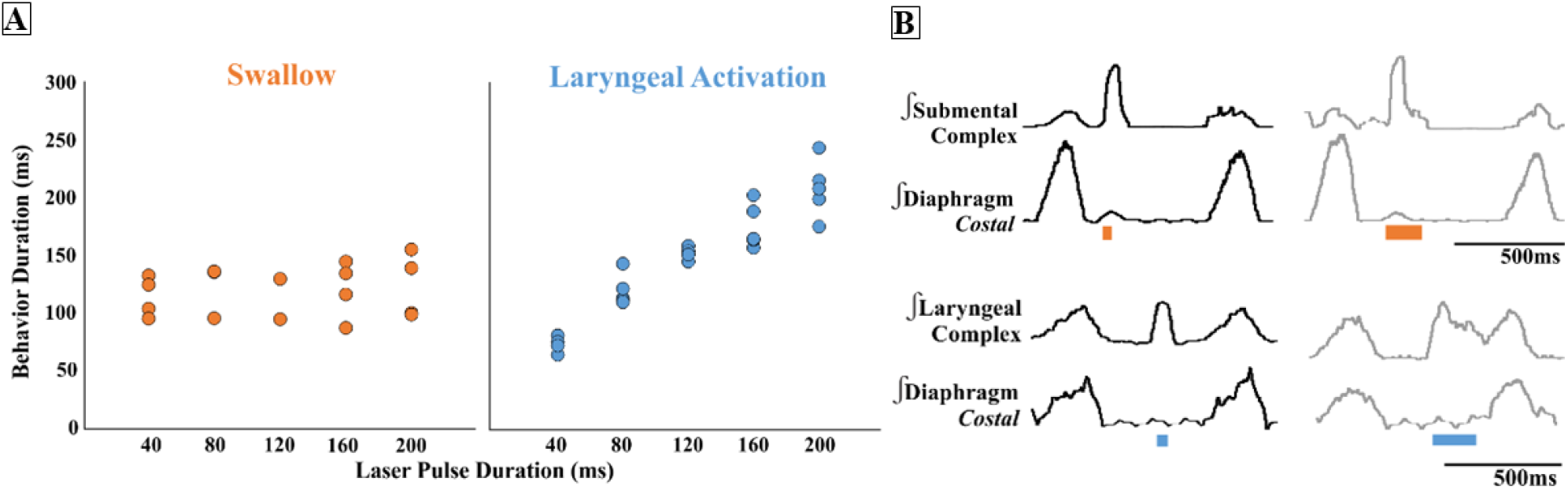
PiCo stimulation triggers swallow behavior whiles activates laryngeal activity. A) Scatter plot of behavior duration versus laser pulse duration for swallow (orange) and laryngeal activation (blue) in ChAT/Vglut2 mice. Each dot represents the average duration per mouse. B) Representative traces of swallow duration shown by submental complex EMG triggered by 40ms pulse in orange on the left and 200ms pulse on the right. Representative traces of laryngeal activation duration shown by laryngeal complex EMG triggered by 40ms pulse in blue on the left and 200ms pulse on the right.

### Swallow related characteristics in water triggered swallows and PiCo triggered swallows

A repeated measures two-way ANOVA revealed no significant differences in swallow onset relative to inspiratory onset between centrally evoked and water evoked swallow (Fig. 4B). A repeated measures two-way ANOVA also revealed PiCo triggered swallows do not occur at a significantly different time than water evoked swallows (Fig. 4C).

**Figure 4.**
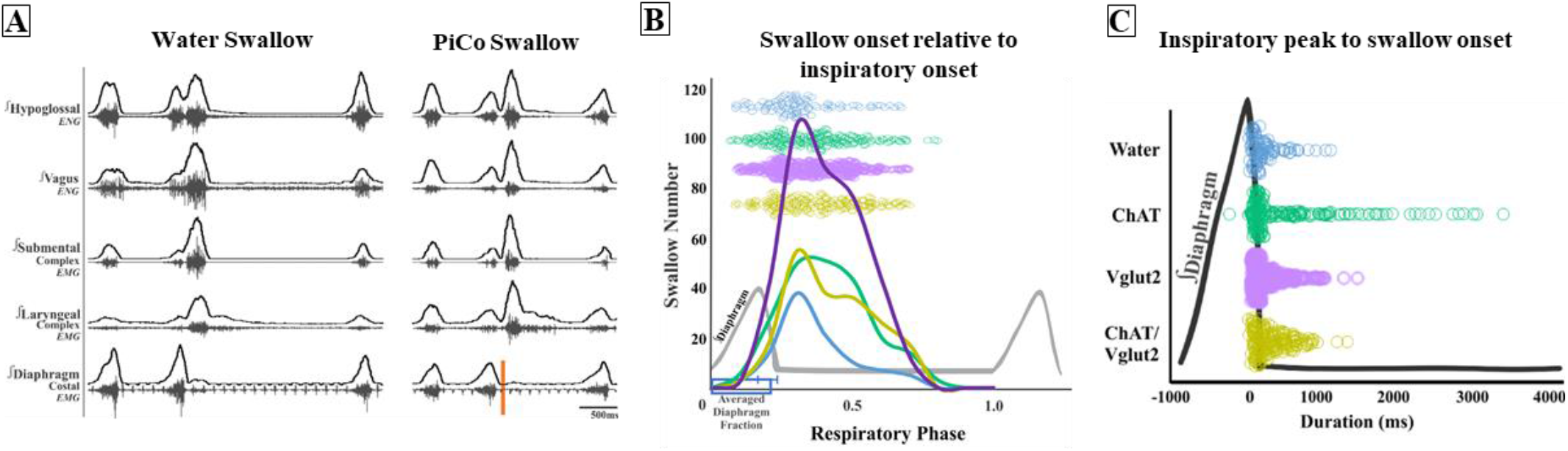
Swallow related characteristics in water triggered swallows and PiCo triggered swallows. A) Representative trace of a swallow triggered by injection of water into the mouth on the left and PiCo stimulation (orange) on the right. B). Histogram of swallows in relation to the onset of inspiration for water swallows (blue, n = 97), ChAT (green, n = 215), Vglut2 (purple, n = 369) and ChAT/Vglut2 (gold, n = 193). C) Dot plot of each swallow in relation to the inspiratory peak. Swallows triggered by water (blue) or PiCo activation occurred at the same time in relation to inspiratory peak.

#### Optogenetic stimulation of ChAT neurons within PiCo

Swallows triggered by ChAT stimulation at the level of PiCo have a trend in decrease duration than swallows evoked by water (290± 125ms) (198 ± 125ms) (*p*= 0.06). There is a significant decrease in XII (297± 129ms) (212 ± 127ms) (*p*= 0.03) and laryngeal complex duration (297 ± 78ms), (163 ± 63ms) (*p*= 0.009). There is significant increase in inspiratory delay (482 ± 696ms) (811 ± 534ms) (*p*= 0.04) in ChAT stimulated swallows. There is no change in swallow motor sequence between swallows evoked by water or ChAT. ChAT stimulated swallows have a significant decrease in XII (73 ± 15) (48 ± 19) % of max (*p*= 0.002) and submental (80 ± 17) (43 ± 40) % of max (*p*= 0.04) amplitude (Fig. S1, Table S2A).

Further comparing male versus female mice, laryngeal complex duration in ChAT stimulated swallows was significantly shorter in female mice (204± 46ms) (109 ± 32ms) (*p*=0.03) (Table S4).

#### Optogenetic stimulation of Vglut2 neurons within PiCo

Swallows triggered by Vglut2 stimulation at the level of PiCo are significantly shorter in duration than swallows evoked by water (256 ± 108ms) (175 ± 94ms) (*p*= 0.006). There is also a significant decrease in XII (286± 95ms) (193 ± 69ms) (*p*= 0.001) and X (256± 76ms) (203 ± 66ms) (*p*= 0.04) nerve duration. Unlike swallows triggered by ChAT stimulation, there is no significant change in diaphragm inter-burst interval or inspiratory delay. Similar to ChAT stimulated swallows, there is no change in swallow motor sequence between swallows evoked by water or Vglut2. Vglut2 stimulated swallows have a significant decrease in X nerve (84 ±13) (58 ± 18) % of max (*p*= 0.003) and submental (88 ± 10) (52 ± 33) % of max (*p*= 0.004) amplitude (Fig. S1, Table S2B).

Further comparing male versus female mice, swallow related inspiratory delay in Vglut2 stimulated swallows was significantly longer in female mice (273± 140ms) (569 ± 256ms) (*p*=0.03) (Table S5).

#### Optogenetic stimulation of ChAT/Vglut2 neurons within PiCo

Five ChAT/Vglut2 mice were stimulated while only 4 triggered swallows and one only stimulated laryngeal activation. There was no significant decrease in swallow related durations triggered by ChAT/Vglut2 stimulation at the level of PiCo most likely due to a low N number. ChAT/Vglut2 stimulated swallows have a significant decrease in X nerve (84 ± 3), (54 ± 5) % of max (*p*= 0.02) and submental (86 ± 16), (49 ± 34) % of max (*p*= 0.05) amplitude (Fig. S1, Table S2C).

There were no sex-specific differences in ChAT/Vglut2 mice. This is most likely due to the low female n number in this group (Table S6).

### Missed or low transfection of PiCo neurons stimulates incomplete swallow related motor activation

Post-hoc histological analysis was performed in the double conditioned cre/FlpO (ChAT/Vglut2) mouse to check the transfection of PiCo neurons after injection of the pAAV-hSyn Con/Fon hChR2(H134R)-EYFP vector (Figs. 5 and 6). Nucleus ambiguous (NA) cholinergic neurons had no transfection and the rostrocaudal distribution of the transgene-expressing neurons was analyzed as represented in the figure 5. In the 5 mice where PiCo stimulation resulted in swallow and/or laryngeal activation response, we found 123 ± 11 neurons expressed EYFP (Fig. 5). Four of the mice that stimulated both swallow and laryngeal activation had an average of 141 transfected neurons while the one mouse that only stimulated laryngeal activation had 46 transfected neurons.

**Figure 5.**
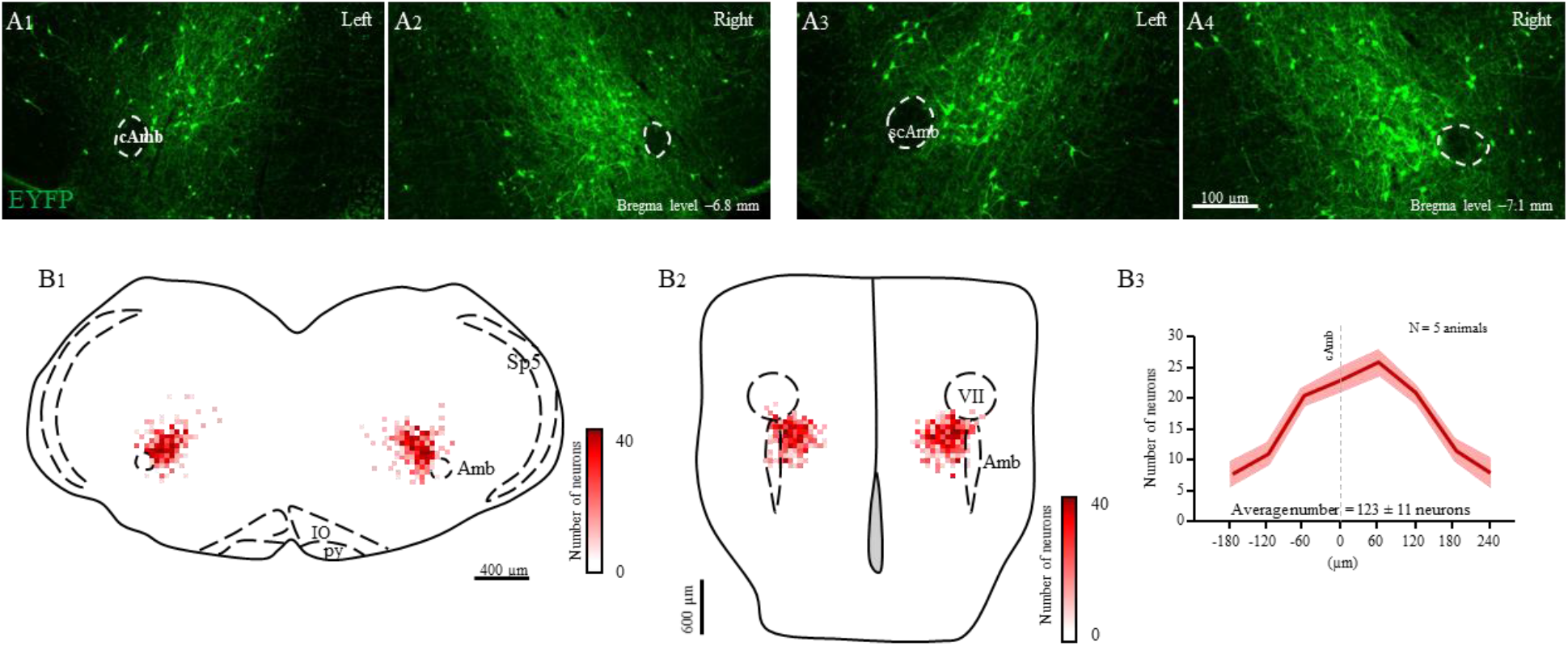
Selective transfection of cholinergic/glutamatergic neurons in PiCo in Chat/Vglut2 mice. A) Transverse hemisection through two different Bregma levels (−6.8 and −7.1 mm) of the transfected neurons into PiCo bilaterally with the pAAV-hSyn Con/Fon hChR2(H134R)-EYFP vector. B) Heat map showing the density of neurons transfected by the pAAV-hSyn Con/Fon hChR2(H134R)-EYFP vector from 1) coronal and 2) ventral view of the 5 animals used in the functional experiments. B3) Rostro-caudal distribution of the total number of transfected neurons counted 1:2 series of 25 μm sections into PiCo. Abbreviations: cAmb, nucleus ambiguus pars compacta; scAmb, nucleus ambiguus pars semi-compacta; IO, inferior olive; py, pyramidal tract; Sp5, spinal trigeminal nucleus; VII, facial motor nucleus.

**Figure 6.**
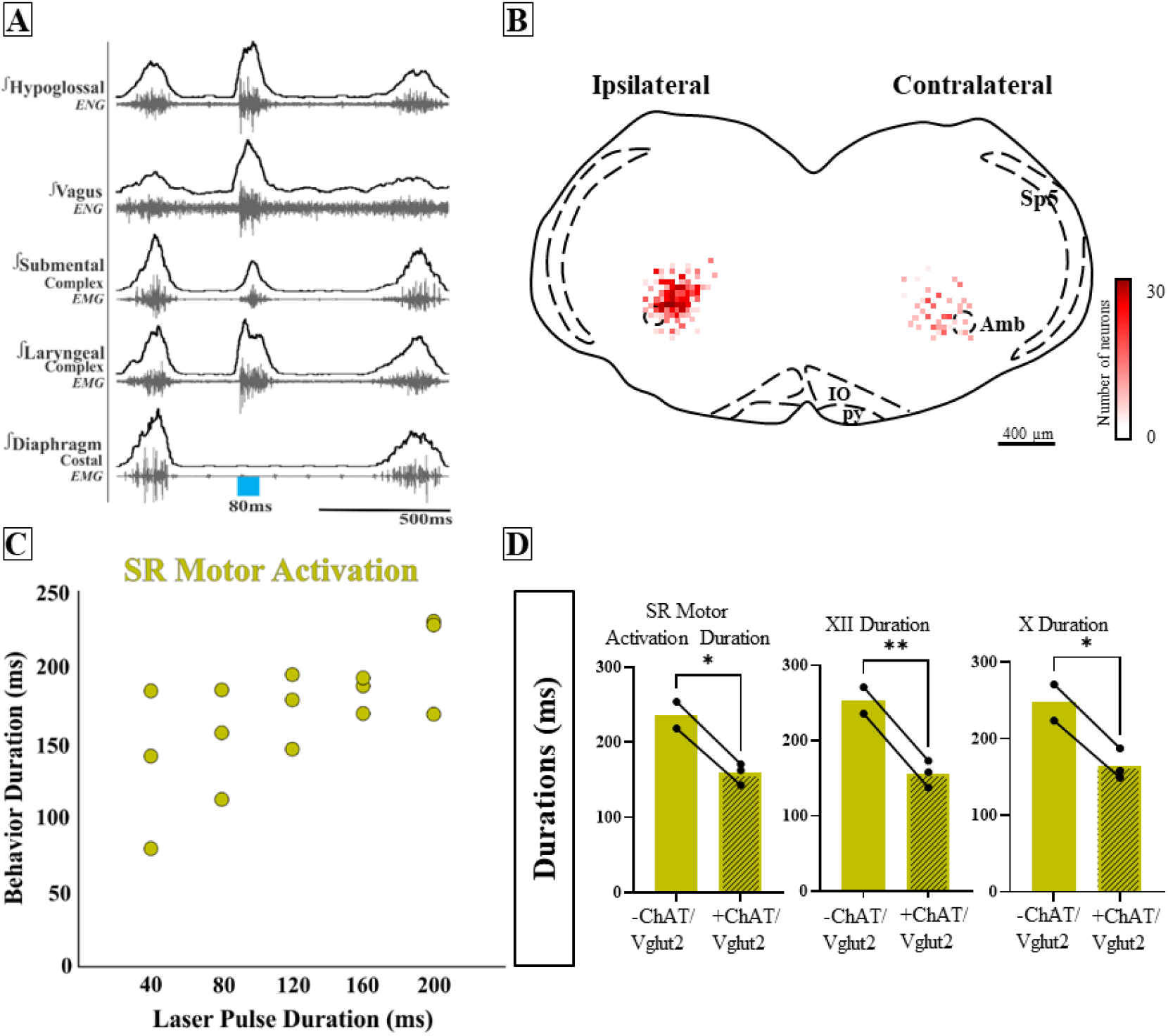
Missed or low transfection of PiCo neurons stimulates incomplete swallow related motor activation. A) Representative trace of 80ms activation of ChAT/Vglut2 neurons at PiCo resulting in swallow related motor activation. B) Heat map showing the density of neurons transfected by the pAAV-hSyn Con/Fon hChR2(H134R)-EYFP vector from coronal view of the 3 animals. Though bilateral transfection, ipsilateral represents the side of the brainstem with the greatest amount of transfection 69 ± 4 neurons and contralateral 26 ± 5 neurons. C) Scatter plot of behavior duration versus laser pulse duration for swallow related motor activation. D) Comparison of total motor activation, hypoglossal (XII) and vagus (X) durations for water swallows (-) and swallow related motor activation in ChAT/Vglut2 mice (N=3). Abbreviations: Amb, nucleus ambiguus; IO, inferior olive; py, pyramidal tract; Sp5, spinal trigeminal nucleus; VII, facial motor nucleus.

However, there were 3 mice that when activated by PiCo, swallow nor laryngeal activation occurred, instead a different behavior was evoked. Across all phases of breathing, activation of PiCo specific neurons evoked a behavior that has similar characteristics to both swallow and laryngeal activation, resulting in a blurred swallow related motor activity response. Swallow motor sequence was reversed with laryngeal activity occurring before submental activity. When compared to a water triggered swallow, swallow related motor activation has a decrease in behavior duration (197 ± 67ms) (159 ± 14ms) (*p*= 0.06), XII duration (131 ± 29ms) (156 ± 18ms) (*p*= 0.003), and X duration (238 ± 69ms) (165 ± 20ms) (*p*= 0.04) (Fig 6, Table S3). There were no sex-specific differences in ChAT/Vglut2 mice. This is most likely due to the low male and female N number in this group (Table S7).

Swallow related motor activation was not triggered, rather stimulated, behavior duration depended on laser pulse duration. Post-hoc histological analysis revealed not only a decrease in total transfection, 95 ± 10 compared to the above 5 mice, but asymmetric transfection. Though bilateral injection, ipsilateral, indicating the side of the brainstem with the most transfection, had on average 69 ± 4 neurons, and the contralateral 26 ± 5 neurons (Fig. 6).

### Characterization of PiCo neurons and projections to swallow and laryngeal related medullary areas

The anatomical description of PiCo region in mice was first described by Anderson, et al. (Anderson et al., 2016). Here, we characterized the distribution of ChAT+ expressing neurons in a coronal segment and heat map of PiCo area (Fig. S2A and B, N = 4 animals). In the rostro-caudal distribution, we found 403 ± 39 ChAT+ neurons, with the most rostral portion of PiCo neurons located next to the caudal pole of the facial nucleus, extending from −6.6 to −7.3 mm distant from Bregma level, reaching caudally to the NA non-compact portion. Also, we can define PiCo neurons being located slightly medial to the NA extending 500 μm medial, and 600 μm dorsal to the NA in a 45° angle.

As described by Anderson et al., ChAT cells in PiCo are also glutamatergic. To characterize this subcluster of neurons we used a triple conditioned mice expressing cre in ChAT cells and FlpO in Vglut2 cells enhanced by a red fluorescent protein (tdTomato, Ai65) inserted into the ROSA26 locus (N = 4 animals). Similar to the previous ChAT staining, the rostro-caudal distribution showed 242 ± 12 neurons ChAT-cre/Vglut2-FlpO Ai65, also represented by a heat map showing the rostro-caudal and medial-lateral distribution (Fig. S2C and D).

To characterize the neuronal inputs from PiCo region to the most known swallow related area (i.e. cNTS), we performed CTb injections into the cNTS (Fig. S3A) and investigated retrograde stained neurons in PiCo area. We found few neurons CTb-labeled in PiCo area (Fig. S3B), showing those areas are slightly connected to produce the swallow behavior as demonstrated in our results. When then performed the CTb injections into the Larynx, targeting the laryngeal complex functionally recorded in this study, in order to investigate a possible direct connection from PiCo to the airway muscle, we did not find retrograde CTb-stained neuron in PiCo area (Fig. S3C).

## Discussion

In the present study we characterized the neuronal coordination of laryngeal motor activity, swallowing and breathing by optogenetically stimulating excitatory neurons in PiCo, a region implicated in the control of postinspiratory behaviors (Anderson et al., 2016). Using transsectional genetics we were able to stimulate specifically interneurons that co-express cholinergic and glutamatergic transmitters and compare their effects with those evoked by stimulating cholinergic and glutamatergic neurons. Taken together, we find that stimulating PiCo glutamatergic/cholinergic neurons triggered swallow motor activity in a phase-dependent and all-or-none manner. Swallowing activity was preferentially evoked when stimulating PiCo during inspiration. Swallow motor activity outlasted the triggering stimulus and either abruptly terminated ongoing inspiratory activity or if the stimulus did not terminate inspiration, swallowing occurred after the completion of inspiratory activity. Thus, PiCo stimulation during inspiration, i.e. when the preBötC is maximally activated, evokes the swallow motor pattern with variable inspiratory inhibition suggesting tight coordination with the respiratory rhythmogenic network (Fig. 7). As already demonstrated by Anderson et al., PiCo is mutually connected with the inspiratory CPG, the preBötC (Anderson et al., 2016). Stimulating Dbx1 neurons in the area of the preBötC inhibits ongoing activity in PiCo, while stimulating PiCo cholinergic neurons inhibits ongoing inspiratory activity and resets respiratory activity *in vitro* and *in vivo*. Here we show that PiCo inhibits inspiration while gating swallow and laryngeal activation (Fig. 7).

**Figure 7.**
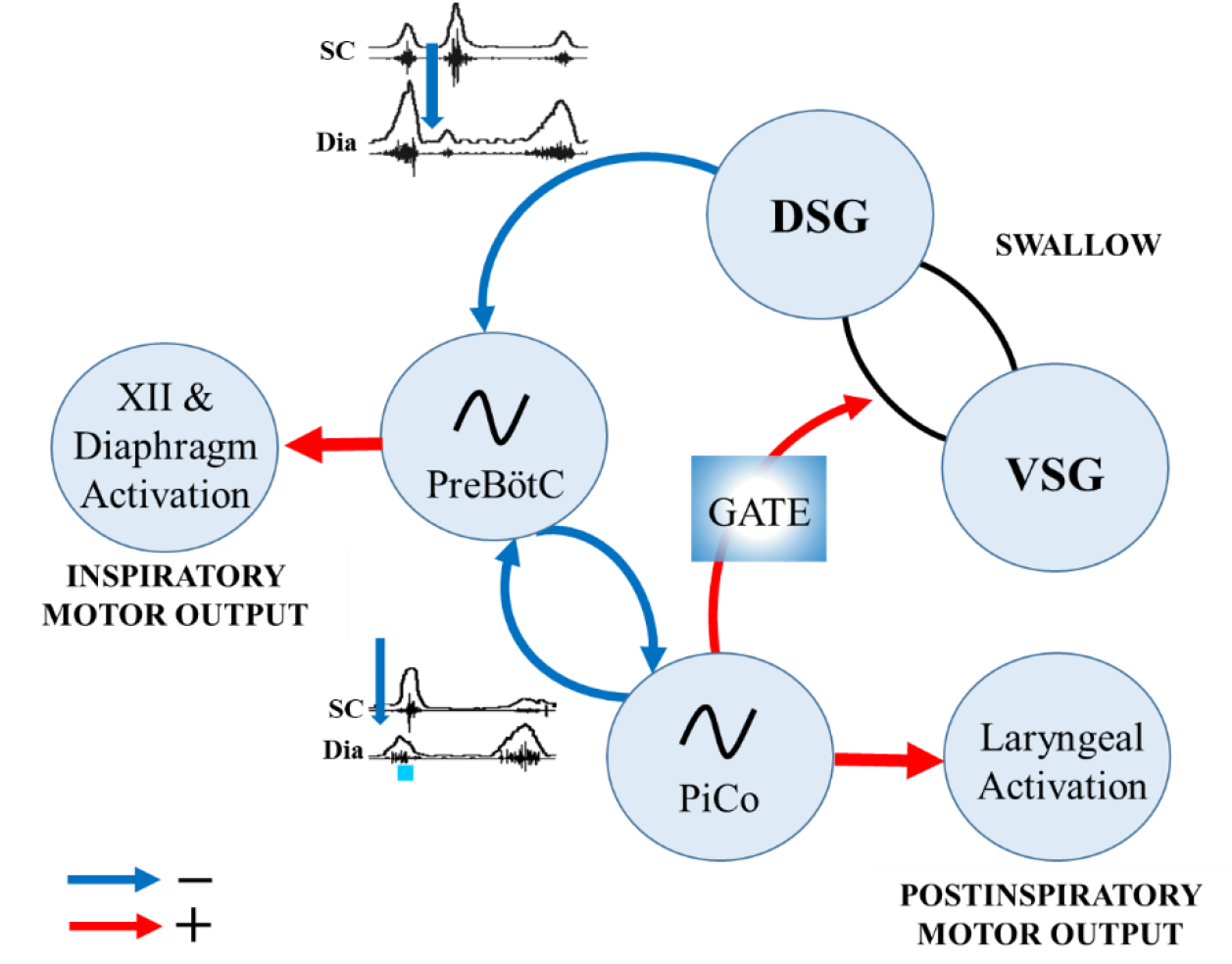
Proposed mechanisms of PiCo interaction with swallow pattern generator (SPG) and inspiratory rhythm generator, preBötC. Anderson et al. previously demonstrated mutual connection between PiCo and preBötC (Anderson et al., 2016). This study further demonstrates inhibitory connections (blue) between SPG and preBötC, and PiCo and preBötC due to the ability for swallow to inhibit inspiration whether evoked by water (Huff et al., 2022) or central PiCo stimulation. PiCo acts as a gating mechanism for postinspiratory behaviors such as swallow and laryngeal activation with excitatory connections (red) to SPG and laryngeal motor neurons.

Just like the gating mechanisms in locomotion (Grillner, 2006), stimulation of PiCo activates different postinspiratory airway protective mechanisms depending on the presence of inspiratory activity (inspiration). Interestingly, for gating to occur, bilateral stimulation of a sufficient number of PiCo neurons is necessary, since incomplete and/or unilateral expression of channelrhodopsin in the population of glutamatergic/cholinergic PiCo neurons led to uncoordinated swallowing attempts characterized by abnormal activation and coordination between laryngeal motoneurons and the submental complex.

While swallow activity was triggered in an all-or-none manner, laryngeal activation was gradually activated for the duration of the optogenetic stimulus (Fig. 3)(Miller & Sherrington, 1915). As shown by Anderson et al. 2016, this laryngeal activity is critical for the generation of postinspiratory activity recorded from the vagal nerve. However, the glutamatergic cholinergic neurons within PiCo are not functioning as laryngeal motor neurons since viral tracing experiments revealed no direct projections to the laryngeal muscles (Fig. S3C). While PiCo activates laryngeal motor activity during postinspiratory activity, the functional role of this motor activity cannot be determined from our recordings. It is conceivable that laryngeal activation via PiCo stimulation could be a central component and integral to a laryngeal adductor reflex (LAR), an airway protective mechanism that prevents aspiration of foreign material (Kaneoka et al., 2018). Another possibility is a muscular response of the, previously studied, non-respiratory expiratory laryngeal motoneurons (Sun, Bautista, Berkowitz, Zhao, & Pilowsky, 2011). Sun et al. suggest the superior laryngeal nerve (SLN) induced expiratory laryngeal motoneuron burst are unable to inhibit inspiratory burst, but a simultaneous signal is sent to a group of neurons that inhibits inspiration for this behavior to occur (Sun et al., 2011).

Our study confirms aspects of Toor et al. who previously hypothesized that neurons of the IRt in the rat act as a hub for swallow and laryngeal activity (Ain Summan Toor et al., 2019). The current study suggests that the same population of glutamatergic/cholinergic neurons can trigger motor patterns of swallow and laryngeal activation, and which of these motor activities is activated by these interneurons is gated by respiratory activity (Fig. 1). It is conceivable that the gating mechanism involves the Parabrachial Nucleus and Kölliker-Fuse Nucleus. Located in the dorsolateral pons this region has been implicated as a sensory relay for the larynx, in particular postinspiratory activity of laryngeal adductor and swallow-breathing coordination (Bautista & Dutschmann, 2014; Dutschmann & Herbert, 2006). Further studies are necessary to understand the interaction between PiCo and the pontine respiratory group on gating swallow and other airway protective behaviors.

Swallow duration and amplitude of swallow related muscles and nerves decreased during PiCo stimulated swallows compared to water evoked swallows (Fig. S1). Swallow motor duration (R. W. Doty & Bosma, 1956) and interneurons in the dorsal swallow group (Yamamoto et al., 2022) respond similarly to fictive SLN stimulation and to physiological water-evoked swallows, though both involve sensory feedback mechanisms. However, when separating out pharyngeal versus laryngeal inputs, sensory-motor coordination during the pharyngeal phase of physiological swallows is more complex (Yamamoto et al., 2022). Effects of pontine respiratory group inhibition on swallow function differed between SLN stimulated swallow and physiological swallows which has been attributed to the sensory feedback of the SPG (Takemura et al., 2022). Though, fictive swallows via SLN stimulation, similar to physiological water-evoked swallows, occur during respiratory phase transitions including late expiration to inspiration and inspiration to postinspiration (Dick, Oku, Romaniuk, & Cherniack, 1993). Physiologic swallows are more dynamic and alter their timing, duration and amplitude to accommodate for change in bolus size, texture and consistency via afferent sensory feedback mechanisms (Dantas et al., 1990; Hrycyshyn & Basmajian, 1972). PiCo triggered swallows preserve the rostro-caudal swallow pattern also seen in physiologic swallows (R. W. Doty & Bosma, 1956; Thexton, Crompton, & German, 2007), though PiCo triggered swallows occur at a broader range of the respiratory cycle, while predominately occurring within the postinspiratory phase (Fig. 4). It is likely that PiCo triggered swallows are not activating the sensory component of the SPG to the same extent as the water evoked swallows, given that the swallows are not associated with upper airway sensory stimulation.

Swallow has been thought of as an “all or nothing” response as early as 1883 (Meltzer, 1883). Whether modulating spinal or vagal feedback (Huff A, 2020b), central drive for swallow/breathing (Huff, Karlen-Amarante, Pitts, & Ramirez, 2022) or lesions in swallow related areas of the brainstem (Car, 1979; Robert W Doty, Richmond, & Storey, 1967; Wang & Bieger, 1991) swallow either occurred or did not. Swallows are thought to be a fixed action pattern, with duration of stimulation having no effect on behavior duration (Fig. 3) (Dick et al., 1993). Thus, it was particularly interesting that in instances when few PiCo neurons were transfected, either unilateral or bilateral, partial swallow related motor activation occurred. Motor activity no longer outlasted laser stimulation rather was contained within, and the timing of swallow related motor sequence was reversed (Fig. 6). Thus, if insufficient numbers of neurons are activated, PiCo’s influence as a gate for airway protective behaviors is blurred, resulting in the uncoordinated activation of muscles involved in both behaviors. This brings into question whether this is the first evidence against the classic dogma of swallow as an “all or nothing” behavior, and/or whether this is an indication that activating the cholinergic/glutamatergic neurons in PiCo is not only gating the SPG, but is actually involved in assembling the swallow motor pattern itself. An incomplete activation of PiCo activates the muscular components of the swallow, without establishing the coordinated timing and sequence of the pattern. The swallow pattern generators (SPG) are thought to consist of bilateral circuits (hemi-CPGs) that govern ipsilateral motor activities, but receive crossing inputs from contralateral swallow interneurons in the reticular formation, thought to coordinate synchrony of swallow movements (Kinoshita et al., 2021; Sugimoto, Umezaki, Takagi, Narikawa, & Shin, 1998; Sugiyama et al., 2011). It is possible that unilateral stimulation of PiCo either desynchronizes swallow interneurons or activates only one side of the SPG. Since we did not record bilateral swallow related muscles and nerves this question needs to be further examined.

Kleinfeld and colleagues introduce the concept of pre-premotor regions “pre^2^motor” to describe that respiratory oscillators can modulate other orofacial premotor oscillators such as whisking and sniffing (McElvain et al., 2018; Moore et al., 2013; Moore et al., 2014) and recently described swallow (Huff et al., 2022; Pitts et al., 2021). By modulating the preBötzinger complex, swallow can be shifted to different times of the respiratory cycle as well as changing swallow related amplitude, laryngeal duration and motor pattern sequence (Huff et al., 2022). Triggering swallow via PiCo activation results in a phase delay of the respiratory cycle, resetting the rhythm, whereas laryngeal activation (non-swallow) has minimal to no effect on respiration phase (Fig. 2). Activation of PiCo specific neurons arrested, or abrogate inspiration triggering swallow, further indicating swallow’s hierarchical control over breathing (fig 2C, 8) (Dick et al., 1993; Huff et al., 2022; Miller & Sherrington, 1915; Pitts et al., 2018).

Diaphragm activity has been shown to be multimodal, having different activity patterns for swallow and breathing, including concurrent inhibited respiratory related activity and activated swallow related activity, *Schluckatmung*, in physiologic and fictive swallows (Huff et al., 2022; Pitts et al., 2021; Pitts et al., 2018). Activation of glutamatergic neurons in the area of PiCo resulted in swallow related diaphragmatic activation, *Schluckatmung*. It has been hypothesized that the SPG activates pre-motor neurons in the dorsal respiratory group responsible for diaphragm recruitment during swallow (Pitts et al., 2018). Inspiratory neurons of the medial reticular formation have been shown to increase firing frequency during *Schluckatmung* (Pitts et al., 2021).

We conclude PiCo acts as a gate for laryngeal coordination during swallow and other behaviors. The identification of PiCo as an important region in swallow-breathing coordination will also be critical for better understanding the mechanisms underlying diseases and disorders with prevalent swallow-breathing discoordination. Leigh Syndrome, Stroke and Parkinson Disease, as well as obstructive sleep apnea and chronic obstructive pulmonary disease all have high incidences of aspiration pneumonia (Armstrong & Mosher, 2011; Cvejic & Bardin, 2018; Su et al., 2014; Tanaka et al., 2022; Won, Byun, Oh, Park, & Seo, 2021). Aspiration is the result of a discoordination of laryngeal closure during swallow that allows foreign material to enter into the airway instead of the esophagus. Further investigation into PiCo in the context of various breathing and neurological diseases can lead to potential targets and therapeutics for decreasing or even eliminating aspiration related pneumonia in high risk populations.

## Methods

### Animals

Adult (P54-131, average P75) both male and female mice were bred at Seattle Children’s Research Institute (SCRI) and used for all experiments. *Vglut2-ires-cre* (*Vglut2)* and *ChAT-ires-cre* (*ChAT*) homozygous breeder lines were obtained from Jackson laboratories (stock numbers 028863 and 031661, respectively). Cre mice were crossed with homozygous mice containing a floxed STOP channelrhodopsin fused to an EYFP (Ai32) reporter sequence from Jackson laboratories (stock number 024109). *ChAT-ires-cre*, and *Vglut2-ires2-FlpO-D*, *technically known as* 129S-Slc17a6^tm1.1(flpo)Hze^/J was obtained from Jackson Laboratories (#031661 and #030212, respectively). To generate double-transgenic mice, the *ChAT* and Vglut2^FlpO^ strains were interbred to generate compound homozygotes, named as ChAT^cre^ Vglut2^FlpO (+/+)^, which tagged neurons that have a developmental history of expressing both *ChAT* and *Vglut2*. Mice were randomly selected from the resulting litters by the investigators. Offspring were group housed with ad libitum access to food and water in a temperature controlled (22 +1 °C) facility with a 12h light/dark cycle. All experiments and animal procedures were approved by the Seattle Children’s Research Institute’s Animal Care and Use Committee and were conducted in accordance with the National Institutes of Health guidelines.

### Brainstem injection of AAV

We restricted ChR2 expression to the PiCo region in order to transfect and photo-stimulate the region with the highest density of ChAT/Vglut2 neurons of the PiCo region (Anderson et al., 2016). For AVV injection, the mice were anesthetized with isoflurane (2%). The correct plane of anesthesia was assessed by the absence of the corneal and hind-paw withdrawal reflexes. Mice received postoperative ketoprofen [7 mg/kg, subcutaneous (s.c.)] for two consecutive days. All surgical procedures were performed under aseptic conditions. The hair over the skull and neck were removed and skin disinfected. The mice were then placed prone on a stereotaxic apparatus (bite bar set at −3.5 mm for flat skull; David Kopf Instruments Tujunga, CA, USA). A 0.5 mm diameter hole was drilled into the occipital plate on the both side caudal to the parieto-occipital suture. Viral solutions were loaded into a 1.2 mm internal diameter glass pipette broken to a 20 μm tip (external diameter). To target the PiCo region with ChR2, the pipette was inserted in the brainstem in the following coordinates: 4.8 mm below the dorsal surface of the cerebellum, 1.1 mm lateral to the midline and 1.6 mm caudal to the lambda and bilateral injections of 150 nL were made, using a glass micropipette and an automatic nanoliter injector (NanoinjectII, Drummond Scientific Co. Broomall, PA).

The mouse strain containing ires-cre and ires-FlpO in ChAT^+^ and Vglut2^+^ respectively, had the PiCo neurons successfully transfected by pAAV-hSyn Con/Fon hChR2(H134R)-EYFP adenovirus vector (cat# 55645-AAV8; AddGene, USA; abbreviated as AAV8-ConFon-ChR2-EYFP) herein named ChAT/Vglut2 in this study. This AAV is a Cre-on/FlpO-on ChR2-EYFP under the synapsin promoter and encoded the photoactivatable cation channel channelrhodopsin-2 (ChR2, H134R) fused to EYFP. The vector was diluted to a final titer of 1 × 10^13^ viral particles/ml with sterile phosphate-buffered saline.

### CTb Injections

Cholera toxin subunit B (CTb) was injected into the cNTS and larynx in three and two mice, respectively. All mice were anesthetized with isoflurane (2%) in 100% oxygen as described previously, while injections of CTb 1% (low salt, 1%, List Biological Laboratories, Campbell, CA) in distilled water, were made using a glass micropipette (tip diameter 15-20 μm) and Nanoinject II, to retrogradely label neurons that innervate these regions. The coordinates to reach cNTS were: 4.2 mm below the dorsal surface of the cerebellum, 0 mm lateral to the midline and 7.8 mm caudal to the Bregma (Kirkcaldie, Watson, Paxinos, & Franklin, 2012). Laryngeal injections were made using the same methods for placement of laryngeal complex electrodes. Seven to ten days following the CTb injections, the mice anesthetized with isoflurane (5%) in 100% oxygen and immediately perfused transcardially with fixative.

### *In Vivo* Experiments

The same experimental protocol was performed for all *Vglut2* and *ChAT* Ai32, as well as *ChAT/Vglut2* mice. Adult mice were initially anesthetized with 100% O_2_ and 1.5% Isoflurane (Aspen Veterinary Resources Ltd, Liberty, MO, USA) for 2-3 minutes in an induction chamber. Once the breathing slowed, they were injected with Urethane (1.5mg.kg, i.p. Sigma Aldrich, St. Louis, MO, USA) and placed supine on a custom surgical table. Core temperature was maintained through a water heating system (PolyScience, Niles, IL, USA) built into the surgical table. Mice were then allowed to spontaneously breathe 100% O2 for the remainder of the surgery and experimental protocol. Adequate depth of anesthesia was determined via heart and breathing rate, as well as lack of toe pinch response every 15 minutes. Bipolar electromyograms (EMG) electrodes in the costal diaphragm were placed to monitor respiratory rate and heart rate throughout the experiment. The trachea was exposed through a midline incision and cannulated caudal to the larynx with a curved (180 degree) tracheal tube (PTFE 24 G, Component Supply, Sparta, TN, USA). The hypoglossal (XII) and vagus (X) nerves were then dissected followed by cannulation of the trachea. The recurrent laryngeal nerve (RLN) was carefully dissected away from each side of the trachea before the cannula was tied in and sealed with super glue to ensure no damage to the RLN. The trachea and esophagus were then cut to detach at the rostral end just caudal to the cricoid cartilage, preserving the arytenoids and bilateral recurrent laryngeal nerves. A tube filled with 100% O_2_ was attached to the cannulated trachea to provide supplemental oxygen throughout the experiment. The occipital bone was removed, followed by continuous perfusion of the medullary surface with warmed (~36°C) artificial cerebral spinal fluid (aCSF; in mM: 118 NaCl, 3 KCl, 25 NaHCO_3_, 1 NaH_2_PO_4_, 1 MgCl_2_, 1.5 CaCl_2_, 30 D-glucose) equilibrated with carbogen (95% O_2_, 5% CO_2_) by a peristaltic pump (Dynamax RP-1, Rainin Instrument Co; Emeryville CA, USA). As previously published (figure 6a (Huff et al., 2022)), the XII and X nerves were isolated unilaterally, cut distally, and recorded from using a fire-polished pulled borosilicate glass (B150-86-15, Sutter Instrument; Novato, CA, USA) filled with aCSF connected to the monopolar suction electrode (A-M Systems, Sequim, WA, USA) and held in a 3D micromanipulator (Narishige, Tokyo, Japan). Multiple bipolar EMGs using 0.002” and 0.003” coated stainless steel wires (A-M Systems, Sequim, WA, USA, part no.790600 and 79100 respectively) according to the techniques of Basmajian and Stecko (Basmajian & Stecko, 1962) simultaneously recorded activity from several swallow- and respiratory-related muscle sites and were placed using hypodermic needles 30G (part no 305106, BD Precision Glide ™, Franklin Lakes, NJ, USA) in the: 1) *submental complex*, which consists of the geniohyoid, mylohyoid and digastric muscles, to determine swallow activity. 2) The *laryngeal complex*,consisting of the arytenoid muscles (transverse, oblique, thyroarytenoid and posterior cricoarytenoid muscles), to determine laryngeal activity during swallows, as well as postinspiratory activity. 3) The *costal diaphragm*, used to measure the multifunctional activity for both inspiration, as well as *Schluckatmung*, a less common diaphragmatic activation during swallow activity. Glass fiber optic (200 um diameter) connected to a blue (447 nm) laser and DPSS driver (Opto Engine LLC, Salt Lake City, Utah, USA) was placed bilaterally in light contact with the brainstem overtop of the predetermined postinspiratory complex (PiCo) (Anderson et al., 2016).

### Stimulation protocols

1) Swallow was stimulated by injecting 0.1cc of water into the mouth using a 1.0cc syringe connected to a polyethylene tube. 2) 25 pulses of each 40ms, 80ms, 120ms, 160ms and 200ms continuous TTL laser stimulation at PiCo was repeated, at random, throughout the respiratory cycle. The lasers were each set to 0.75mW and triggered using Spike2 software (Cambridge Electronic Design, Cambridge, UK). These stimulation protocols were performed in all *Vglut2, ChAT* Ai32 mice as well as *ChAT/Vglut2* mice.

### Analysis

All electroneurogram (ENG) and EMG activity were amplified and band-pass filtered (0.03 – 1 KHz) by a differential AC Amplifier (A-M System model 1700, Sequim, WA, USA), acquired in a A/D converter (CED 1401; Cambridge Electronic Design, Cambridge, UK) then integrated, rectified, smoothed and stored using Spike2 software (Cambridge Electronic Design, Cambridge, UK).

We evaluated swallows that were trigged by injection of water into the mouth as well as behaviors in response to laser stimulation at PiCo: swallow, laryngeal activation and no response. Swallow was characterized as a delayed response to the laser outlasting the laser duration, activation of XII, X, submental and laryngeal complex, and a submental-laryngeal delay. Diaphragm activity during PiCo triggered swallows (*schluckatmung*) was present in some animals but this was not common. Laryngeal activation was characterized as activity of the XII, X, laryngeal complex from onset to offset of the laser pulse, and absence of the diaphragm. The submental complex was active in some animals but not all during laryngeal activation. No response was characterized as lack of response to the laser and was grouped with laryngeal activation for the non-swallow analysis in respiratory phase shift plots (Fig. 2). *Swallow duration* was determined by the onset to the termination of the submental complex. In the case the submental complex muscles were not available then it was determined by the onset to the offset of the XII. *Swallow sequence* was calculated as the time difference between the peak of the laryngeal and submental complex. *Schluckatmung* duration was determined by the onset to the offset of the diaphragm during a swallow. *Laryngeal activation duration* was determined by the onset to the termination of the laryngeal complex. *Diaphragm inter-burst interval* was calculated as the offset of the diaphragm to the onset of the proceeding breathing. *Inspiratory delay* was calculated as the offset of the swallow related laryngeal activity to the onset of the proceeding breath. Duration of each nerve and muscle was determined by the onset to the offset of that respective nerve/muscle during swallow.

As previously reported (figure 6d (Huff et al., 2022)). Respiratory phase reset curves calculated by defining the respiratory cycle as the onset of the diaphragm to the onset of the subsequent diaphragm activity. The *phase shift* elicited by each stimulation of water was calculated as the duration of the respiratory cycle containing the stimulus, divided by the preceding respiratory cycle. The phase of the swallow stimulation (*respiratory phase*) was calculated as the time between the onset of the inspiration (diaphragm) and the stimulus onset, divided by the expected phase. The average phase shift was then plotted against the respiratory phase in bins containing 1/10 of the expected phase (Baertsch, Baertsch, & Ramirez, 2018). Swallow histogram plots were created by the phase of breathing in which swallow occurred in, calculated as the onset of inspiration to the onset of swallow divided by the respiratory cycle duration and plotted against the number of swallows that occurred within the 1/10 binned respiratory phase. Swallow was also plotted in relation to the peak activation of the diaphragm as a duration with zero equaling the peak of the inspiratory related diaphragm activity.

Probability plots were calculated by assigning a “0” to the no response behavior or a “0 or 1” to the laryngeal activation or swallow behavior. These number were then averaged and plotted against the *respiratory phase* and binned to 1/10 of the respiratory phase.

All data are expressed as mean ± standard deviation (SD). Statistical analyses were performed using GraphPad Prism 9 (GraphPad Software®, Inc. La Jolla, USA). For comparison between baseline and stimulus within the same group was made by paired Student t-test. For comparison between different genetic lines in the probability plots two-way ANOVA was used. Differences were considered significant at *p* ≥ 0.05. Investigators were not blinded during analysis. Sample sizes were chosen on the basis of previous studies.

### Histology

At the end of experiments, all animals were deeply anesthetized with 5% isoflurane in 100% oxygen and perfused through the ascending aorta with 20 ml of phosphate buffered saline (PB; pH 7.4) followed by 4% phosphate-buffered (0.1 M; pH 7.4; 20 ml) paraformaldehyde (Electron Microscopy Sciences, Fort Washington, PA). The brains were removed and stored in the perfusion fixative for 4 h at 4 °C, followed by 20% sucrose for 8h. Series of coronal sections (25 μm) from the brains were cut using a cryostat and stored in cryoprotectant solution at −20°C (20% glycerol plus 30% ethylene glycol in 50 ml phosphate buffer, pH 7.4) prior to histological processing. All histochemical procedures were done using free-floating sections.

*ChAT* was detected using a polyclonal goat anti-ChAT antibody (AB144P; Millipore; 1:100), CTb was detected using a monoclonal mouse anti-CTb (AB62429; ABCAM; 1:5000) and EYFP was detected using a polyclonal mouse anti-GFP (06-896, Millipore; 1:1000) diluted in PB containing 2% normal donkey serum (017-000-121, Jackson Immuno Research Laboratories) and 0.3% Triton X-100 and incubated for 24 h. Sections were subsequently rinsed in PB and incubated for 2 h in an Alexa 488 donkey anti-goat antibody (711-545-152; 1:250; Jackson Immuno Research Laboratories), Alexa 594 donkey anti-mouse antibody (715-585-150; 1:400; Jackson Immuno Research Labratories) or Alexa 647 donkey anti-mouse (A31571; 1:400; Life technologies). For all secondary antibodies used, control experiments confirmed that no labeling was observed when primary antibodies were omitted. The sections were mounted on slides in rostrocaudal sequential order, dried, and covered with fluoromount (00-4958-02; Thermo Fisher). Coverslips were affixed with nail polish.

Sections were also examined to confirm the transfected cells. As shown in Figure 5 and S2, according to the Paxinos and Franklin mouse atlas (Kirkcaldie et al., 2012), the transfected cells were located just dorsal to the Nucleus ambiguus near Bregma level −6.84 mm, ~1100 μm from the midline, and ~700 μm above the marginal layer.

### Cell counting, Imaging and Data analysis

A VS120-S6-W Virtual Slide Scanner (Olympus) was used to scan all the sections. Images were taken with a color camera (Nikon DS-Fi3). To restrict any influences on our counted results, the photomicrography and counting were performed by one blind researcher. Image J (version 1.41; National Institutes of Health, Bethesda, MD) was used for cell counting and Canvas software (ACD Systems, Victoria, Canada, v. 9.0) was used for line drawings. A one-in-two series of 25-μm brain sections was used per mouse, which means that each section analyzed was 50 μm apart. The area analyzed was delimited based on previously reports (Anderson et al., 2016) (mean of 5,423 μm^2^). The sections were counted bilaterally, averaged and the numbers reported as mean ± SEM. Section alignment were relative to a reference section, as previously described (Anderson et al., 2016) and based on Paxinos and Franklin (Kirkcaldie et al., 2012) (2012).

## Acknowledgements

We are grateful for NIH grants P01 HL090554 (awarded to J.M.R.), R01 HL144801 (awarded to J. M.R.), R01 HL 151389 (awarded to J.M.R.), F32 HL160102-01 (awarded to A.H.) for funding this project.

## Data Availability

All data is publicly available (10.6084/m9.figshare.21909819).

---

**Supplemental Figure 1.**
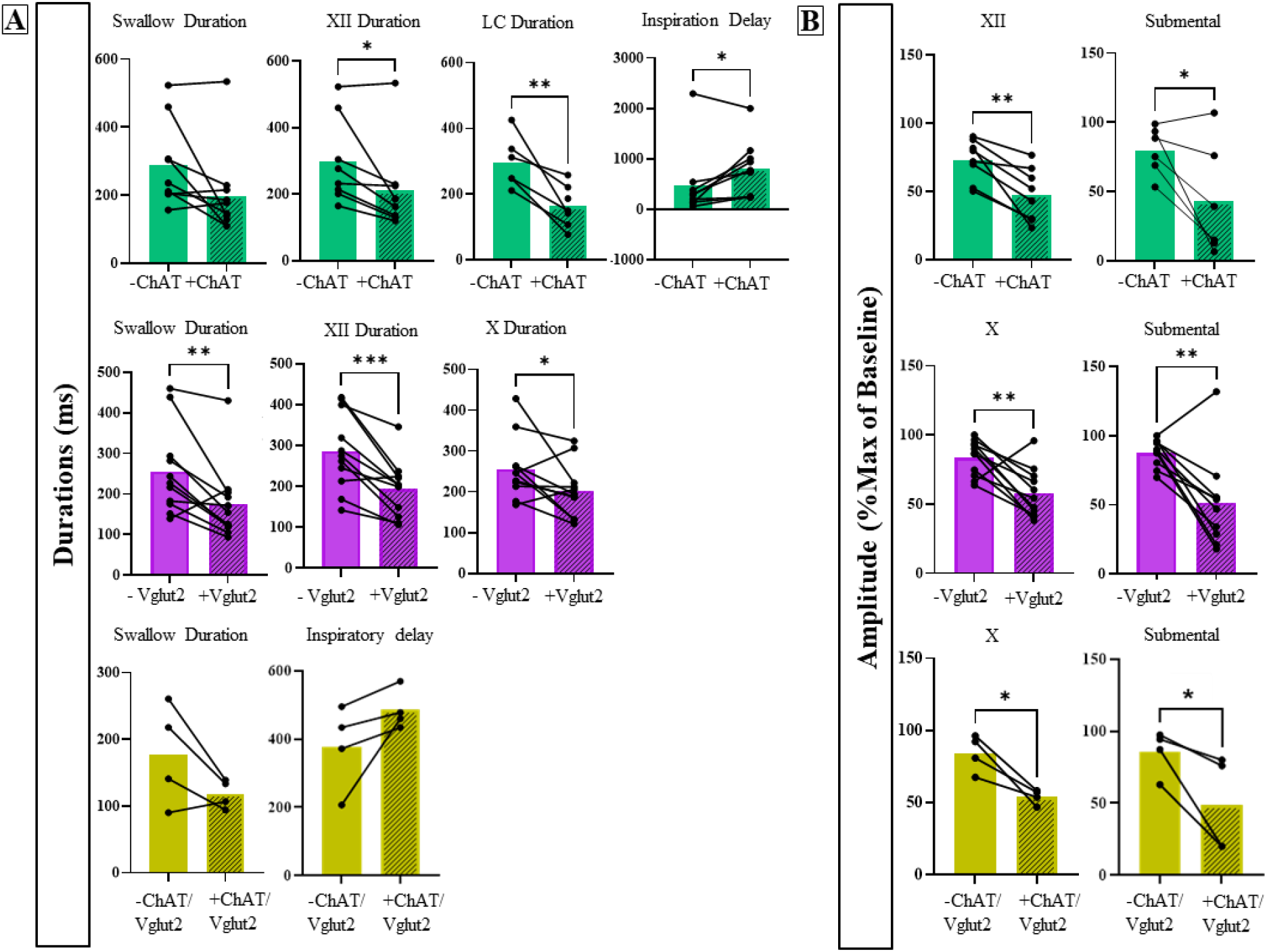
PiCo triggered swallows have a decrease in duration and amplitude compared to water triggered swallows. A) Comparison of durations and B) amplitude in swallow-related characteristics for water swallows (-) and PiCo stimulated swallows (+) in ChAT (green, N=10), Vglut2 (purple, N=11) and ChAT/Vglut2 (gold, N=4). Abbreviations: X, vagus nerve; XII, hypoglossal nerve; LC, laryngeal complex.

**Supplemental Figure 2.**
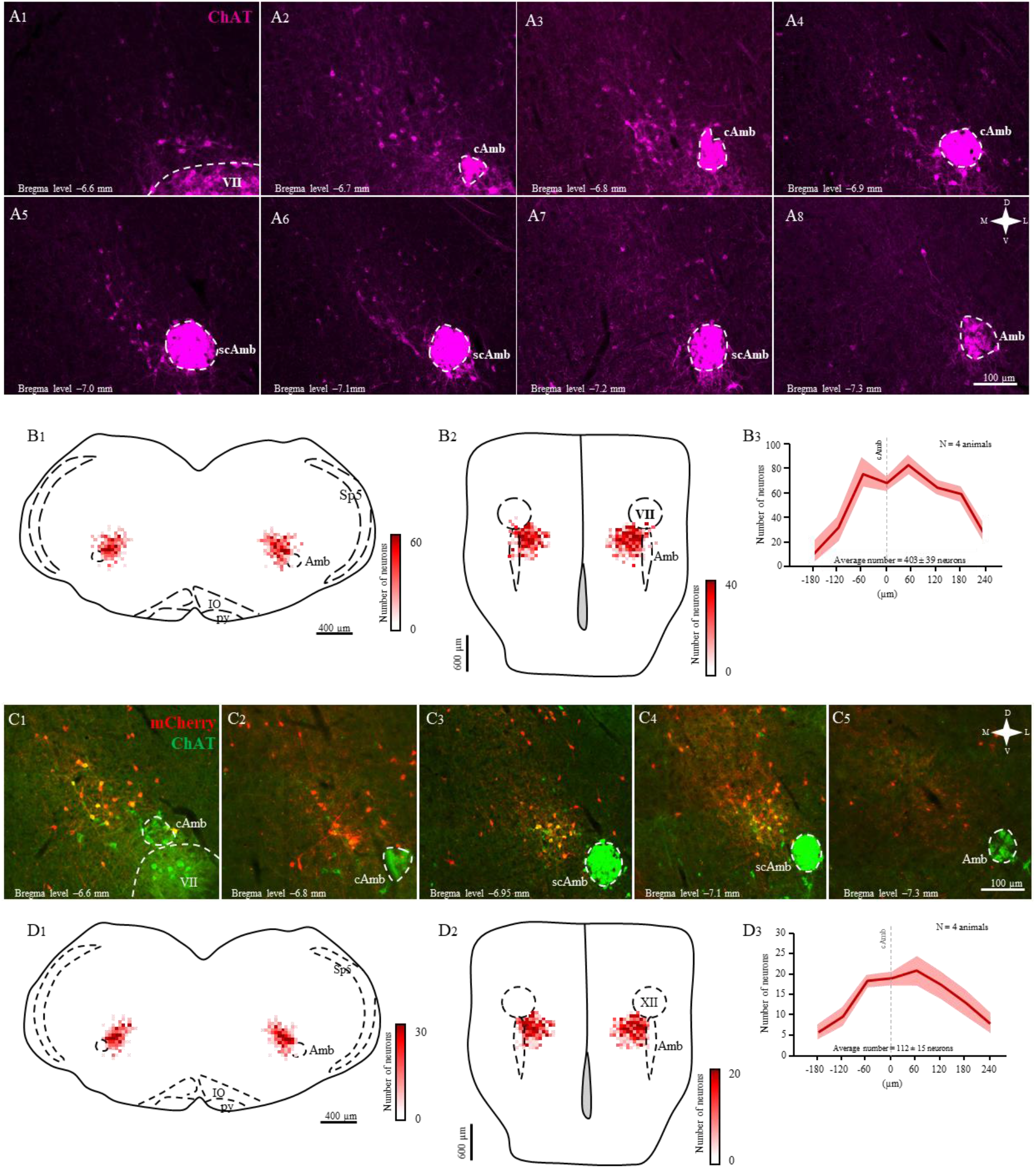
Anatomical characterization of PiCo region. A) Coronal views (Bregma level −6.6 to −7.3 mm) of the ventro-medial medulla showing the location of the ChAT neurons (magenta) in PiCo region. B) Heat map showing the density of ChAT immunoreactive neurons from 1) coronal and 2) ventral view of 4 animals. B3) Rostro-caudal distribution of the total number of ChAT immunoreactive counted 1:2 series of 25 μm sections into PiCo. C) Coronal views (Bregma level −6.6 to −7.3 mm) of the ventro-medial medulla showing the location of the double conditioned ChAT/Vglut2/Ai65 neurons (red) in PiCo region. D) Heat map showing the density of ChAT/Vglut2/Ai65 neurons from 1) coronal and 2) ventral view of 4 animals. D3) Rostro-caudal distribution of the total number of ChAT/Vglut2/Ai65 neurons counted 1:2 series of 25 μm sections into PiCo. Abbreviations: cAmb, nucleus ambiguus pars compacta; scAmb, nucleus ambiguus pars semi-compacta; Amb, nucleus ambiguus pars non-compacta; VII, facial motor nucleus; IO, inferior olive; py, pyramidal tract; Sp5, spinal trigeminal nucleus; VII, facial motor nucleus.

**Supplemental Figure 3.**
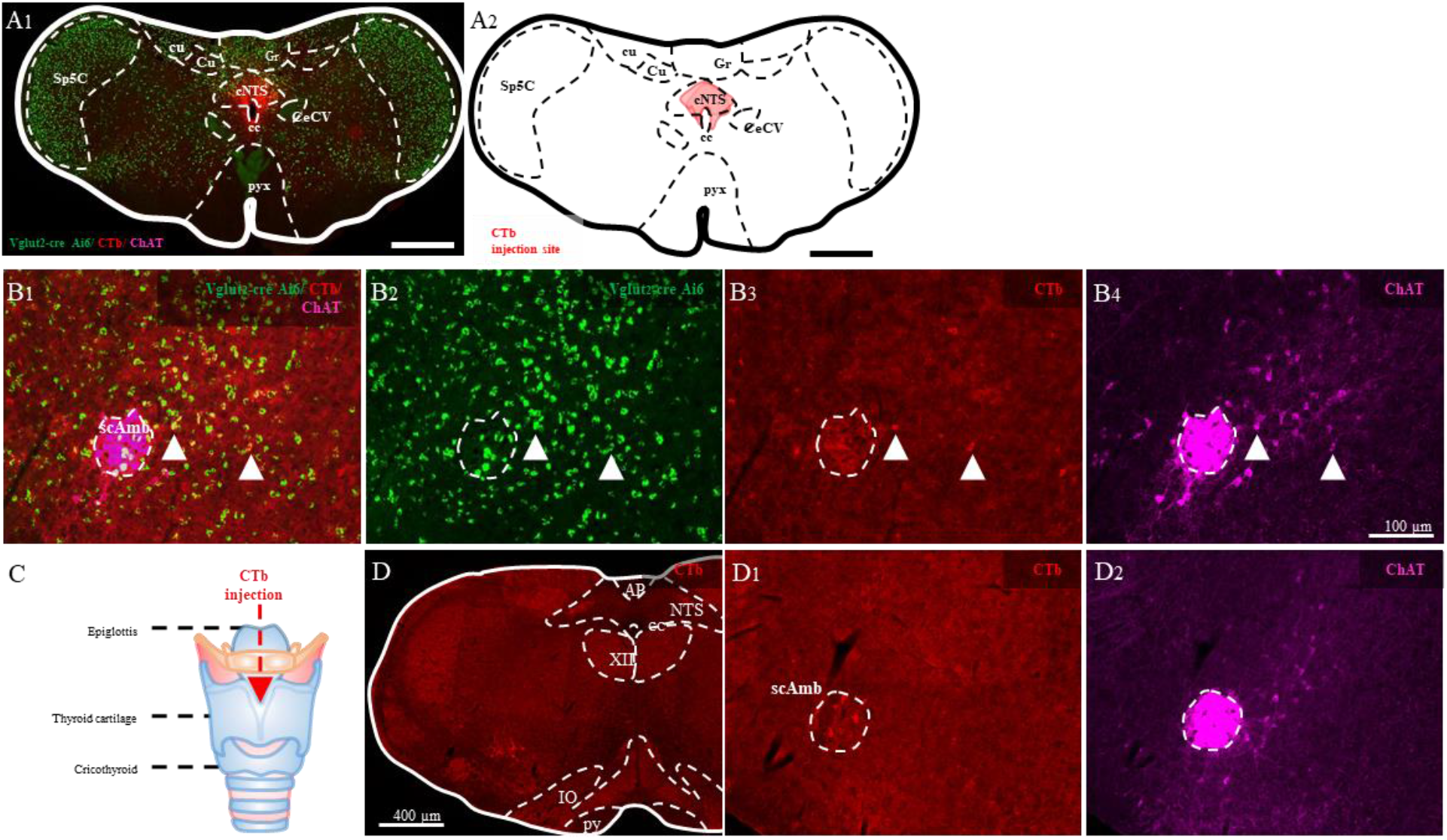
PiCo cells retrogradely labelled with CTb following deposit into cNTS. A) Photomicrography and schematic drawing of retrograde tracer CTb injected into cNTS. B) Coronal views (Bregma level −7.1 mm) of PiCo showing the location of the retrogradely stained cells with CTb (red), ChAT (magenta) and Vglut2 (green) neurons in PiCo region. White arrows indicate some examples of triple retrogradely labelled cells in PiCo. C) Drawing of the larynx where CTb has been deposited. D) Coronal views (Bregma level −7.1 mm) of PiCo showing no CTb (red) stained neurons after CTb injection into the larynx. Abbreviations: scAmb, nucleus ambiguus pars semi-compacta; Sp5C, caudal part of spinal trigeminal nucleus; Cu, cuneate nucleus; cu, cuneate fasciculus; AP, area postrema; Gr, gracile nucleus; cc, central canal; CeCV, central cervical nucleus; cNTS, caudal nucleus of solitary tract; IO, inferior olive; py, pyramidal tract; pyx, pyramidal decussation; XII, hypoglossal nucleus.

**Table S1.** Provides means and standard deviations (SD) for vagus and laryngeal complex duration during laryngeal activation in response to increasing stimuli in ChAT, Vglut2, and ChAT/Vglut2 mice.

| **Laryngeal Activation** | **+ ChAT Stimulation** | **+ Vglut2 Stimulation** | **+ ChAT/Vglut2 Stimulation** |
| --- | --- | --- | --- |
|  | **mean ( SD )** | **mean ( SD )** | **mean ( SD )** |
| **40ms** |  |  |  |
| Vagus Duration (ms) | 77 ( 14 ) | 85 ( 8 ) | 75 ( 11 ) |
| Laryngeal Complex Duration (ms) | 76 ( 9 ) | 81 ( 13 ) | 72 ( 7 ) |
| **80ms** |  |  |  |
| Vagus Duration (ms) | 108 ( 16 ) | 101 ( 27 ) | 106 ( 12 ) |
| Laryngeal Complex Duration (ms) | 105 ( 16 ) | 103 ( 23 ) | 119 ( 14 ) |
| **120ms** |  |  |  |
| Vagus Duration (ms) | 140 ( 31 ) | 123 ( 55 ) | 142 ( 23 ) |
| Laryngeal Complex Duration (ms) | 131 ( 20 ) | 125 ( 57 ) | 151 ( 6 ) |
| **160ms** |  |  |  |
| Vagus Duration (ms) | 163 ( 25 ) | 114 ( 62 ) | 142 ( 42 ) |
| Laryngeal Complex Duration (ms) | 164 ( 32 ) | 128 ( 68 ) | 175 ( 19 ) |
| **200ms** |  |  |  |
| Vagus Duration (ms) | 197 ( 62 ) | 132 ( 50 ) | 165 ( 61 ) |
| Laryngeal Complex Duration (ms) | 243 ( 146 ) | 152 ( 57 ) | 208 ( 25 ) |
| **Average** |  |  |  |
| Vagus Duration (ms) | 145 ( 33 ) | 100 ( 24 ) | 127 ( 30 ) |
| Laryngeal Complex Duration (ms) | 133 ( 40 ) | 103 ( 28 ) | 149 ( 13 ) |

**Table S2.** Provides means, standard deviations (SD), *p*-values and the direction of change for swallow related parameters when evoked by water (water swallows) and optogenetic stimulation of PiCo in A) ChAT, B) Vglut2 and C) ChAT/Vglut2 mice.

| A | Water Swallow |  | + ChAT Stimulation |  |  |  |
| --- | --- | --- | --- | --- | --- | --- |
|  | mean | ( SD ) | mean | ( SD ) | p -value | Change |
| (n=10) |  |  |  |  |  |  |
| Swallow Duration (ms) | 290 | ( 125 ) | 198 | ( 125 ) | 0.06 | - |
| Hypoglossal Duration (ms) | 297 | ( 129 ) | 212 | ( 127 ) | 0.03 | ↓ |
| Vagus Duration (ms) | 294 | ( 101 ) | 224 | ( 112 ) | 0.05 | ↓ |
| Laryngeal Complex Duration (ms) | 297 | ( 78 ) | 163 | ( 63 ) | 0.009 | ↓ |
| *Schluckatmung* Duration (ms) | 228 | ( - ) | - | ( - ) | - | - |
| Diaphragm Inter-Burst Interval (ms) | 1002 | ( 691 ) | 1361 | ( 938 ) | 0.06 | - |
| SR Inspiratory Delay (ms) | 482 | ( 696 ) | 811 | ( 534 ) | 0.04 | ↑ |
| Swallow Sequence (ms) | 24 | ( 40 ) | 18 | ( 19 ) | 0.29 | - |
| Swallow Onset (ms) | 268 | ( 133 ) | 433 | ( 472 ) | 0.24 | - |
| Hypoglossal Amplitude (% max) | 73 | ( 15 ) | 48 | ( 19 ) | 0.002 | ↓ |
| Vagus Amplitude (% max) | 73 | ( 17 ) | 53 | ( 27 ) | 0.06 | - |
| Submental Complex Amplitude (% max) | 78 | ( 17 ) | 43 | ( 40 ) | 0.04 | ↓ |
| Laryngeal Complex Amplitude (% max) | 74 | ( 23 ) | 55 | ( 36 ) | 0.07 | - |
| *Schluckatmung* Amplitude (% max) | - | ( - ) | - | ( - ) | - | - |

| B | Water Swallow |  | + Vglut2 Stimulation |  |  |  |
| --- | --- | --- | --- | --- | --- | --- |
|  | mean | ( SD ) | mean | ( SD ) | p -value | Change |
| (n=11) |  |  |  |  |  |  |
| Swallow Duration (ms) | 256 | ( 108 ) | 175 | ( 94 ) | **0.007** | ↓ |
| Hypoglossal Duration (ms) | 286 | ( 95 ) | 193 | ( 69 ) | **0.001** | ↓ |
| Vagus Duration (ms) | 256 | ( 76 ) | 203 | ( 66 ) | **0.04** | ↓ |
| Laryngeal Complex Duration (ms) | 311 | ( 149 ) | 250 | ( 100 ) | 0.20 | - |
| *Schluckatmung* Duration (ms) | 150 | ( 48 ) | 176 | ( 70 ) | 0.83 | - |
| Diaphragm Inter-Burst Interval (ms) | 834 | ( 488 ) | 749 | ( 316 ) | 0.54 | - |
| SR Inspiratory Delay (ms) | 437 | ( 438 ) | 381 | ( 232 ) | 0.68 | - |
| Swallow Sequence (ms) | 29 | ( 38 ) | 22 | ( 30 ) | 0.37 | - |
| Swallow Onset (ms) | 160 | ( 103 ) | 200 | ( 119 ) | 0.40 | - |
| Hypoglossal Amplitude (% max) | 83 | ( 13 ) | 77 | ( 30 ) | 0.38 | - |
| Vagus Amplitude (% max) | 84 | ( 13 ) | 58 | ( 18 ) | **0.003** | ↓ |
| Submental Complex Amplitude (% max) | 88 | ( 10 ) | 52 | ( 33 ) | **0.004** | ↓ |
| Laryngeal ComplexAmplitude (% max) | 78 | ( 16 ) | 85 | ( 76 ) | 0.76 | - |
| *Schluckatmung* Amplitude (% max) | 80 | ( 12 ) | 170 | ( 152 ) | 0.62 | - |

| C | Water Swallow |  | + ChAT/Vglut2 Stimulation |  |  |  |
| --- | --- | --- | --- | --- | --- | --- |
|  | mean | ( SD ) | mean | ( SD ) | p -value | Change |
| (n=4) |  |  |  |  |  |  |
| Swallow Duration (ms) | 178 | ( 76 ) | 119 | ( 21 ) | 0.14 | - |
| Hypoglossal Duration (ms) | 221 | ( 82 ) | 131 | ( 34 ) | 0.15 | - |
| Vagus Duration (ms) | 233 | ( 86 ) | 162 | ( 31 ) | 0.08 | - |
| Laryngeal Complex Duration (ms) | 251 | ( 83 ) | 230 | ( 138 ) | 0.79 | - |
| *Schluckatmung* Duration (ms) | - | ( - ) | - | ( - ) | - | - |
| Diaphragm Inter-Burst Interval (ms) | 746 | ( 80 ) | 866 | ( 306 ) | 0.37 | - |
| SR Inspiratory Delay (ms) | 377 | ( 124 ) | 486 | ( 59 ) | 0.11 | - |
| Swallow Sequence (ms) | 55 | ( 41 ) | 50 | ( 59 ) | 0.84 | - |
| Swallow Onset (ms) | 185 | ( 53 ) | 231 | ( 142 ) | 0.51 | - |
| Hypoglossal Amplitude (% max) | 85 | ( 12 ) | 72 | ( 25 ) | 0.39 | - |
| Vagus Amplitude (% max) | 84 | ( 13 ) | 54 | ( 5 ) | **0.02** | ↓ |
| Submental Complex Amplitude (% max) | 86 | ( 16 ) | 49 | ( 34 ) | **0.05** | ↓ |
| Laryngeal ComplexAmplitude (% max) | 81 | ( 15 ) | 71 | ( 41 ) | 0.52 | - |
| *Schluckatmung* Amplitude (% max) | - | ( - ) | - | ( - ) | - | - |

**Table S3.**
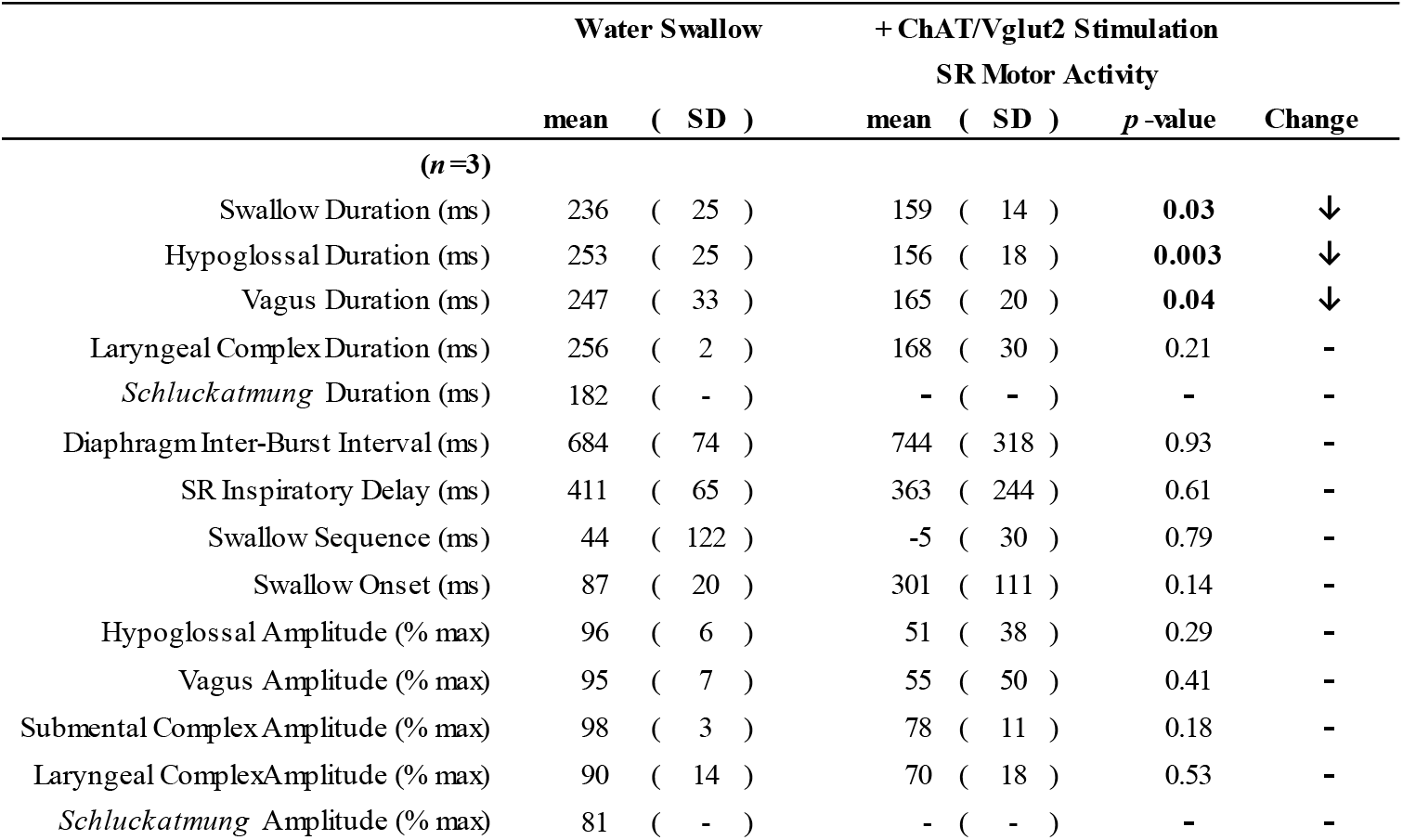
Provides means, standard deviations (SD), *p*-values and the direction of change for swallow related parameters when evoked by water (water swallows) and optogenetic stimulation of PiCo ChAT/Vglut2 mice evoking swallow related motor activity.

|  | Water Swallow |  | + ChAT/Vglut2 Stimulation |  |  |
| --- | --- | --- | --- | --- | --- |
|  | mean | ( SD ) | mean ( SD ) | *p*-value | Change |
| **(*n* = 3)** |  |  |  |  |  |
| Swallow Duration (ms) | 236 | ( 25 ) | 159 ( 14 ) | **0.03** | ↓ |
| Hypoglossal Duration (ms) | 253 | ( 25 ) | 156 ( 18 ) | **0.003** | ↓ |
| Vagus Duration (ms) | 247 | ( 33 ) | 165 ( 20 ) | **0.04** | ↓ |
| Laryngeal Complex Duration (ms) | 256 | ( 2 ) | 168 ( 30 ) | 0.21 | - |
| *Schluckatmung* Duration (ms) | 182 | ( - ) | - ( - ) | - | - |
| Diaphragm Inter-Burst Interval (ms) | 684 | ( 74 ) | 744 ( 318 ) | 0.93 | - |
| SR Inspiratory Delay (ms) | 411 | ( 65 ) | 363 ( 244 ) | 0.61 | - |
| Swallow Sequence (ms) | 44 | ( 122 ) | -5 ( 30 ) | 0.79 | - |
| Swallow Onset (ms) | 87 | ( 20 ) | 301 ( 111 ) | 0.14 | - |
| Hypoglossal Amplitude (% max) | 96 | ( 6 ) | 51 ( 38 ) | 0.29 | - |
| Vagus Amplitude (% max) | 95 | ( 7 ) | 55 ( 50 ) | 0.41 | - |
| Submental Complex Amplitude (% max) | 98 | ( 3 ) | 78 ( 11 ) | 0.18 | - |
| Laryngeal Complex Amplitude (% max) | 90 | ( 14 ) | 70 ( 18 ) | 0.53 | - |
| *Schluckatmung* Amplitude (% max) | 81 | ( - ) | - ( - ) | - | - |

**Table S4.** Provides means, standard deviations (SD), *p*-values and the direction of change for swallow related parameters between male and female mice during water swallows and PiCo stimulated swallows in ChAT mice.

| Water Swallow | Male (*n*=5) |  | Female (*n*=4) |  | *p*-value | Change |
| --- | --- | --- | --- | --- | --- | --- |
|  | mean | ( SD ) | mean | ( SD ) |  |  |
| Swallow Duration (ms) | 295 | ( 134 ) | 284 | ( 133 ) | 0.90 | - |
| XII Duration (ms) | 308 | ( 127 ) | 280 | ( 158 ) | 0.79 | - |
| Vagus Duration (ms) | 294 | ( 107 ) | 292 | ( 115 ) | 0.98 | - |
| Laryngeal Complex Duration (ms) | 299 | ( 46 ) | 295 | ( 115 ) | 0.95 | - |
| *Schluckatmung* Duration (ms) | 228 | ( - ) | - | ( - ) | - | - |
| Diaphragm Inter-Burst Interval (ms) | 1261 | ( 835 ) | 678 | ( 304 ) | 0.23 | - |
| SR Inspiratory Delay (ms) | 717 | ( 900 ) | 188 | ( 71 ) | 0.29 | - |
| Swallow Sequence (ms) | 1 | ( 5 ) | 53 | ( 47 ) | **0.04** | ↑ |
| Swallow Onset (ms) | 308 | ( 101 ) | 218 | ( 167 ) | 0.35 | - |
| + ChAT Stimulation | Male (*n*=6) |  | Female (*n*=4) |  | *p*-value | Change |
|  | mean | ( SD ) | mean | ( SD ) |  |  |
| Swallow Duration (ms) | 239 | ( 150 ) | 137 | ( 36 ) | 0.23 | - |
| XII Duration (ms) | 244 | ( 147 ) | 147 | ( 35 ) | 0.31 | - |
| Vagus Duration (ms) | 262 | ( 121 ) | 150 | ( 33 ) | 0.17 | - |
| Laryngeal Complex Duration (ms) | 204 | ( 46 ) | 109 | ( 32 ) | **0.03** | ↓ |
| *Schluckatmung* Duration (ms) | - | ( - ) | - | ( - ) | - | - |
| Diaphragm Inter-Burst Interval (ms) | 1510 | ( 985 ) | 1138 | ( 954 ) | 0.57 | - |
| SR Inspiratory Delay (ms) | 921 | ( 584 ) | 646 | ( 477 ) | 0.46 | - |
| Swallow Sequence (ms) | 16 | ( 19 ) | 22 | ( 22 ) | 0.64 | - |
| Swallow Onset (ms) | 429 | ( 522 ) | 438 | ( 461 ) | 0.98 | - |

**Table S5.** Provides means, standard deviations (SD), *p*-values and the direction of change for swallow related parameters between male and female mice during water swallows and PiCo stimulated swallows in Vglut2 mice.

| Water Swallow | Male (*n*=7) |  | Female (*n*=4) |  | *p*-value | Change |
| --- | --- | --- | --- | --- | --- | --- |
|  | mean | ( SD ) | mean | ( SD ) |  |  |
| Swallow Duration (ms) | 254 | ( 108 ) | 258 | ( 125 ) | 0.95 | - |
| XII Duration (ms) | 295 | ( 92 ) | 269 | ( 112 ) | 0.68 | - |
| Vagus Duration (ms) | 248 | ( 59 ) | 270 | ( 110 ) | 0.68 | - |
| Laryngeal Complex Duration (ms) | 307 | ( 106 ) | 317 | ( 227 ) | 0.92 | - |
| *Schluckatmung* Duration (ms) | 158 | ( 52 ) | 120 | ( - ) | - | - |
| Diaphragm Inter-Burst Interval (ms) | 796 | ( 509 ) | 902 | ( 515 ) | 0.75 | - |
| SR Inspiratory Delay (ms) | 423 | ( 541 ) | 461 | ( 229 ) | 0.90 | - |
| Swallow Sequence (ms) | 30 | ( 46 ) | 29 | ( 23 ) | 0.97 | - |
| Swallow Onset (ms) | 141 | ( 86 ) | 195 | ( 136 ) | 0.43 | - |
| + Vglut2 Stimulation | Male (*n*=7) |  | Female (*n*=4) |  | *p*-value | Change |
|  | mean | ( SD ) | mean | ( SD ) |  |  |
| Swallow Duration (ms) | 198 | ( 109 ) | 136 | ( 47 ) | 0.32 | - |
| XII Duration (ms) | 207 | ( 79 ) | 170 | ( 50 ) | 0.42 | - |
| Vagus Duration (ms) | 224 | ( 69 ) | 166 | ( 47 ) | 0.17 | - |
| Laryngeal Complex Duration (ms) | 282 | ( 100 ) | 195 | ( 86 ) | 0.18 | - |
| *Schluckatmung* Duration (ms) | 176 | ( 70 ) | - | ( - ) | - | - |
| Diaphragm Inter-Burst Interval (ms) | 645 | ( 207 ) | 932 | ( 422 ) | 0.16 | - |
| SR Inspiratory Delay (ms) | 273 | ( 140 ) | 569 | ( 256 ) | **0.03** | ↑ |
| Swallow Sequence (ms) | 21 | ( 38 ) | 25 | ( 9 ) | 0.88 | - |
| Swallow Onset (ms) | 174 | ( 110 ) | 247 | ( 135 ) | 0.35 | - |

**Table S6.**
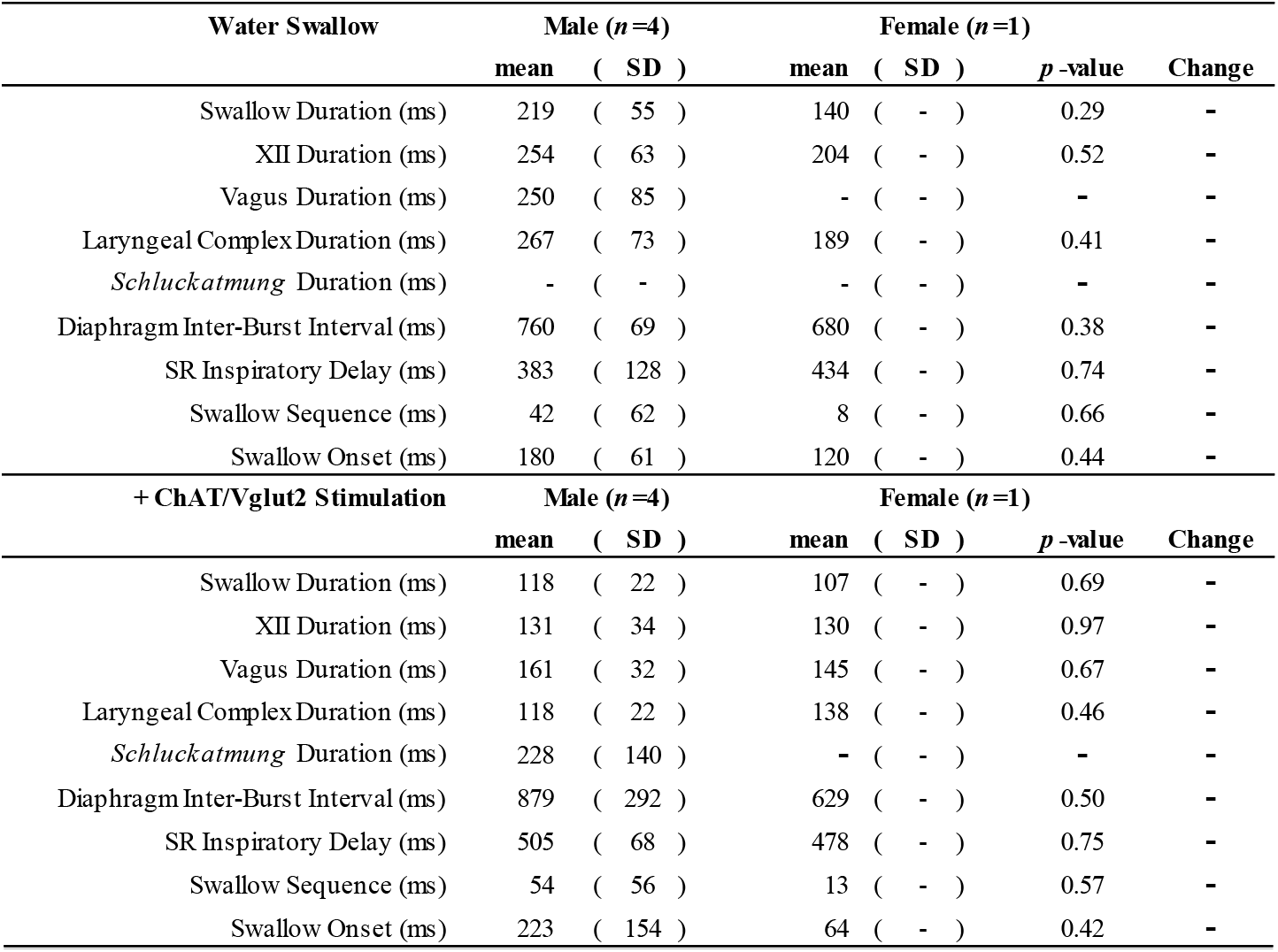
Provides means, standard deviations (SD), *p*-values and the direction of change for swallow related parameters between male and female mice during water swallows and PiCo stimulated swallows in ChAT/Vglut2 mice.

| Water Swallow | Male (*n*=4) |  | Female (*n*=1) |  | *p*-value | Change |
| --- | --- | --- | --- | --- | --- | --- |
|  | mean | ( SD ) | mean | ( SD ) |  |  |
| Swallow Duration (ms) | 219 | ( 55 ) | 140 | ( - ) | 0.29 | - |
| XII Duration (ms) | 254 | ( 63 ) | 204 | ( - ) | 0.52 | - |
| Vagus Duration (ms) | 250 | ( 85 ) | - | ( - ) | - | - |
| Laryngeal ComplexDuration (ms) | 267 | ( 73 ) | 189 | ( - ) | 0.41 | - |
| *Schluckatmung* Duration (ms) | - | ( - ) | - | ( - ) | - | - |
| Diaphragm Inter-Burst Interval (ms) | 760 | ( 69 ) | 680 | ( - ) | 0.38 | - |
| SR Inspiratory Delay (ms) | 383 | ( 128 ) | 434 | ( - ) | 0.74 | - |
| Swallow Sequence (ms) | 42 | ( 62 ) | 8 | ( - ) | 0.66 | - |
| Swallow Onset (ms) | 180 | ( 61 ) | 120 | ( - ) | 0.44 | - |
| + ChAT/Vglut2 Stimulation | Male (*n*=4) |  | Female (*n*=1) |  | *p*-value | Change |
|  | mean | ( SD ) | mean | ( SD ) |  |  |
| Swallow Duration (ms) | 118 | ( 22 ) | 107 | ( - ) | 0.69 | - |
| XII Duration (ms) | 131 | ( 34 ) | 130 | ( - ) | 0.97 | - |
| Vagus Duration (ms) | 161 | ( 32 ) | 145 | ( - ) | 0.67 | - |
| Laryngeal ComplexDuration (ms) | 118 | ( 22 ) | 138 | ( - ) | 0.46 | - |
| *Schluckatmung* Duration (ms) | 228 | ( 140 ) | - | ( - ) | - | - |
| Diaphragm Inter-Burst Interval (ms) | 879 | ( 292 ) | 629 | ( - ) | 0.50 | - |
| SR Inspiratory Delay (ms) | 505 | ( 68 ) | 478 | ( - ) | 0.75 | - |
| Swallow Sequence (ms) | 54 | ( 56 ) | 13 | ( - ) | 0.57 | - |
| Swallow Onset (ms) | 223 | ( 154 ) | 64 | ( - ) | 0.42 | - |

**Table S7.** Provides means, standard deviations (SD), *p*-values and the direction of change for swallow related parameters between male and female mice during water swallows and PiCo stimulated swallow related motor activity in ChAT/Vglut2 mice.

| **Water Swallow** | **Male (n=1)** |  | **Female (n=1)** |  | **p -value** | **Change** |
| --- | --- | --- | --- | --- | --- | --- |
|  | **mean** | **( SD )** | **mean** | **( SD )** |  |  |
| Swallow Duration (ms) | 254 | ( - ) | 218 | ( - ) | - | - |
| XII Duration (ms) | 271 | ( - ) | 236 | ( - ) | - | - |
| Vagus Duration (ms) | 271 | ( - ) | 224 | ( - ) | - | - |
| Laryngeal Complex Duration (ms) | 254 | ( - ) | 257 | ( - ) | - | - |
| *Schluckatmung* Duration (ms) | - | ( - ) | 182 | ( - ) | - | - |
| Diaphragm Inter-Burst Interval (ms) | 736 | ( - ) | 632 | ( - ) | - | - |
| SR Inspiratory Delay (ms) | 457 | ( - ) | 365 | ( - ) | - | - |
| Swallow Sequence (ms) | -42 | ( - ) | 130 | ( - ) | - | - |
| Swallow Onset (ms) | 102 | ( - ) | 73 | ( - ) | - | - |
| **+ ChAT/Vglut2 Stimulation** | **Male (n=1)** |  | **Female (n=2)** |  | **p -value** | **Change** |
| **SR Motor Activity** | **mean** | **( SD )** | **mean** | **( SD )** |  |  |
| Swallow Duration (ms) | 171 | ( - ) | 153 | ( 14 ) | 0.48 | - |
| XII Duration (ms) | 173 | ( - ) | 148 | ( 15 ) | 0.39 | - |
| Vagus Duration (ms) | 187 | ( - ) | 154 | ( 6 ) | 0.14 | - |
| Laryngeal Complex Duration (ms) | 195 | ( - ) | 155 | ( 28 ) | 0.45 | - |
| *Schluckatmung* Duration (ms) | - | ( - ) | - | ( - ) | - | - |
| Diaphragm Inter-Burst Interval (ms) | 1023 | ( - ) | 604 | ( 291 ) | 0.45 | - |
| SR Inspiratory Delay (ms) | 507 | ( - ) | 292 | ( 297 ) | 0.66 | - |
| Swallow Sequence (ms) | 22 | ( - ) | -19 | ( 26 ) | 0.42 | - |
| Swallow Onset (ms) | 423 | ( - ) | 240 | ( 49 ) | 0.20 | - |

